# A disynaptic circuit in the globus pallidus controls locomotion inhibition

**DOI:** 10.1101/2020.07.19.211318

**Authors:** Asier Aristieta, Massimo Barresi, Shiva A. Lindi, Gregory Barriere, Gilles Courtand, Brice de la Crompe, Lise Guilhemsang, Sophie Gauthier, Stéphanie Fioramonti, Jérôme Baufreton, Nicolas P. Mallet

## Abstract

Basal ganglia (BG) inhibit movement through two independent pathways, the indirect- and the hyperdirect-pathways. The globus pallidus (GP) has always been viewed as a simple relay within these two pathways, but its importance has changed drastically with the discovery of two functionally-distinct cell types, namely the prototypic and the arkypallidal neurons. Classic BG models suggest that all GP neurons receive GABAergic inputs from striato-pallidal indirect spiny projection neurons and glutamatergic inputs from subthalamic neurons. However, whether this synaptic connectivity scheme applies to both GP cell-types is currently unknown. Here, we optogenetically dissect the input organization of prototypic and arkypallidal neurons and further define the circuit mechanism underlying action inhibition in BG. Our results highlight that an increased activity of arkypallidal neurons is required to inhibit locomotion. Finally, this work supports the view that arkypallidal neurons are part of a novel disynaptic feedback loop that broadcast inhibitory control on movement execution.

## Introduction

Action execution requires the selection of appropriate motor programs while suppressing unwanted ones and this function is crucial for interaction and survival in an ever-changing environment. The basal ganglia (BG) circuits form highly-conserved neuronal networks in vertebrates (Grillner et al., 2013), and their activity is critical to gate the motor components and adjust the vigor of goal-directed actions while also being involved in many parallel brain functions including motivation, decision-making, and associative learning (Cox and Witten, 2019; Hwang et al., 2019; Matamales et al., 2020; Roseberry et al., 2016; Rueda-orozco and Robbe, 2015; Tecuapetla et al., 2016; Yttri and Dudman, 2016). Rate-coding models of BG functional organization predict that actions are selected or inhibited depending on the level of activity of their GABAergic outputs (Albin et al., 1989; Alexander and Crutcher, 1990). Indeed, the tonic activity of BG output nuclei, exerts sustain inhibition onto thalamus and brainstem motor programs (Cebrián et al., 2005; Hikosaka, 2007; Morrissette et al., 2019; Roseberry et al., 2016; Rossi et al., 2016). Accordingly, action facilitation requires a disinhibition mechanism induced by the suppression of BG output activity, a condition obtained when the so-called direct pathway—the projections from D1-expressing striatal neurons (D1-SPNs) to BG outputs—is activated (Chevalier and Deniau, 1990; Jin and Costa, 2010; Kravitz et al., 2010; Roseberry et al., 2016; Schmidt et al., 2013). In contrast, action inhibition has been associated with increased BG output activity and two BG network mechanisms have been considered so far to underlie such effect. The first requires activation of the indirect pathway (Kravitz et al., 2010; Roseberry et al., 2016). Indeed, activation of this polysynaptic route—that includes projections from D2-expressing striatal neurons (D2-SPNs), the external globus pallidus (GP), and subthalamic nucleus (STN) to BG outputs— leads to an increase in BG output firing and has thus been considered as a ‘No Go’ signal. The second one is dependent upon the activation of the hyperdirect pathway, which provides a fast route linking the cortex to BG outputs through the STN, and it has been associated with a ‘stop’ signal (Pasquereau and Turner, 2017; Schmidt et al., 2013).

Although helpful in providing a general framework to apprehend BG function and dysfunction, BG models have been challenged by many functional and structural findings (Cazorla et al., 2015). In particular, one limiting aspect is the key contribution of the GP in these circuits which has been previously overlooked (Gittis et al., 2014). Indeed, the GP is a hub nucleus at the interface of the indirect and hyperdirect pathways integrating a large amount of striatal D2-SPN and STN inputs, while its long-range GABAergic projections target the whole BG to control and orchestrate BG neuronal activity (Bevan et al., 1998; Crompe et al., 2020; Dodson et al., 2015; Fujiyama et al., 2016; Glajch et al., 2016; Kita, 2007; Mallet et al., 2012, 2016; Mastro et al., 2014, 2017; Sato et al., 2000; Smith et al., 1998). Since D2-SPN and STN inputs are of opposite nature at the level of the GP (i.e. inhibitory for D2-SPN and excitatory for STN inputs), it is not clear how different modulation of GP activity can lead to the same inhibitory effect on motor action. In addition, while classic models assume the GP to be composed of only one cell population projecting only to the STN, recent findings have described an overall dichotomous organization of GP neurons at the molecular, structural and functional level (Abdi et al., 2015; Dodson et al., 2015; Mallet et al., 2012). Prototypic GP neurons represent the main population with ~70% of all GP neurons and have the ‘classic’ features of GP neurons: tonic activity and long-range projections to the substantia nigra pars reticulata (SNR) and the STN (Bevan et al., 1998; Mallet et al., 2012; Mastro et al., 2014). This principal cell population can also be further divided into subpopulations based on molecular markers, projection sites and differential contribution to motor functions (Abdi et al., 2015; Abecassis et al., 2020; Bevan et al., 1998; Dodson et al., 2015; Fujiyama et al., 2016; Mastro et al., 2014, 2017; Saunders et al., 2015). On the contrary, arkypallidal neurons represent a unique cell population reaching ~30% of all GP cells and project massively only to the striatum (Mallet et al., 2012). In addition, in rats performing a stop-signal task, arkypallidal neuron firing can efficiently suppress ongoing striatal activity (Mallet et al., 2016) suggesting the existence of a novel ‘Cancel’ pathway not previously accounted for by BG models and which provides the animal with a mechanism to interrupt imminent actions independently from the traditional indirect ‘No Go’ pathway. This functional GP organization raises important questions regarding how both cell populations integrate striatal D2-SPN and STN inputs. In this work, we used optogenetic approaches combined with *in vivo* and *ex vivo* recordings to dissect the functional connectome of D2-SPN and STN inputs onto identified arkypallidal and prototypic GP neurons. We also assessed the functional contribution of both pathways in the context of locomotion control. Our results show the existence of a novel disynaptic pathway differentially recruited by the indirect and hyperdirect pathways that can efficiently control the animal locomotion. These findings place the GP in a hub position to powerfully switch ongoing actions from execution to inhibition.

## Results

### Identification of prototypic and arkypallidal neurons in D2-Cre and Vglut2-Cre transgenic mice

Prototypic and arkypallidal neurons have been described to present different *in vivo* and *ex vivo* electrical properties in both wild-type animals (Abdi et al., 2015; Dodson et al., 2015; Mallet et al., 2016) and transgenic mice (Abrahao and Lovinger, 2018; Hernandez et al., 2015; Mastro et al., 2014). To confirm the distinct electrophysiological properties of the two identified GP-cell population in transgenic D2-Cre and Vglut2-Cre mice, we recorded GP neuron activity *in vivo* and *ex vivo* and labelled cells with neurobiotin/biocytin for post-hoc immunochemical identification of the recorded neurons (***Figure S1***). Cell-type classification into prototypic and arkypallidal neurons was based on the differential expression profile of specific transcription factors: prototypic neurons expressed the transcription factor Homeobox protein Nkx-2.1 (***Figure S1A***) whereas arkypallidal neurons were immunopositive for forkhead-box P2 (FoxP2) (***Figure S1B***). The electrical activities of Nkx-2.1+ prototypic and FoxP2+ arkypallidal neurons were noticeably different both *in vivo* and *ex vivo*, in agreement with previous reports (Abdi et al., 2015; Dodson et al., 2015). In particular, prototypic neurons had a significantly higher firing rate and were more regular (***Figure S1E-J, supplementary table S7***) than arkypallidal neurons which often had an irregular activity including long epoch of silence, especially *ex vivo* (***Figure S1E-J, supplementary table S7***). These distinct electrical properties were also used to build a larger database that includes all recorded but unlabelled putative prototypic and arkypallidal GP cell-types (***Figure S1E-J***). Comparisons of the data collected from putative GP neurons were in agreement with the data obtained from identified cells (***Figure S1E-J, supplementary table S7***).

### Striatopallidal inputs differentially impact onto prototypic and arkypallidal neurons

The striato-pallidal (i.e. D2-SPNs) projections represent the principal GABAergic inputs to the GP (Kita, 2007). To investigate the functional connectome of striato-pallidal inputs onto prototypic and arkypallidal neurons, we used D2-Cre mice injected with an AAV containing a floxed Channelrhodopsin (AAV-DIO-ChR2) in the striatum (***Figure 1A***). We first validated this D2-Cre∷ AAV-DIO-ChR2 opto-excitatory approach by testing the expression of preproenkephalin in a sample of opto-tagged striatal neurons (***Figure 1B, C***). Application of 2s long blue light pulses induced a strong excitation in all identified D2-SPNs (***Figure 1D***). We then studied the downstream impact of such D2-SPN photo-activation on the spontaneous firing activity of identified GP neurons *in vivo*. We found that prototypic neurons were profoundly inhibited during D2-SPN opto-excitation (***Figure 1B, E***) whereas arkypallidal (***Figure 1B, F***) and STN (***Figure 1B, G***) neurons were both robustly excited. To further dissect the functional dynamics of D2-SPN inputs with finer temporal resolution, we then applied opto-stimulation using light pulses of 10 ms durations (***Figure S2A***). These shorter stimulations evoked more realistic firing activity in D2-SPN neurons, yet it reproduced the core findings obtained with longer stimulation protocols. Indeed, all prototypic neurons exhibited a fast inhibitory response (mean of 8.7 ± 0.4 ms) hinting at a direct monosynaptic connectivity while the recovery of the baseline firing rate was relatively slow (***Figure S2B-D***, ***supplementary table S8***). Interestingly, the excitatory responses induced in arkypallidal and STN neurons parallel nicely the time-course of prototypic inhibition (***Figure S2B-D, supplementary table S8***). Importantly, this opposition of firing rate responses, observed both upon long and short D2-SPNs opto-excitations between prototypic vs. arkypallidal and STN neurons, was identical in our larger dataset including all recorded putative GP cell populations (***Figure S3, supplementary table S9***). This confirms that these stereotypical responses are robust features underlying the functional dynamics that exist between these different cell populations. To further investigate the synaptic mechanism leading to such opposition of *in vivo* firing between GP neurons, we next performed *ex vivo* patch-clamp recordings on prototypic and arkypallidal neurons (***Figure 1H-J, supplementary table S1***) in D2-Cre∷ AAV-DIO-ChR2 and D2-Cre∷ Ai32 mice (***Figure S4A, B***). This latter mouse line unequivocally ensures that all D2-SPNs are expressing ChR2, thus ruling out the possibility that differential neuronal responses in GP is caused by improper viral vector expression. Characterization of D2-SPN inputs using these two optogenetic strategies reveals similar proportion of input-recipient prototypic and arkypallidal neurons (***Figure S4C, D, supplementary table S10)***, therefore data obtained from D2-Cre∷ AAV-ChR2 and D2-Cre∷ Ai32 were pulled together for both identified (***Figure 1K-L)*** and putative ***(Figure S4E, F)*** prototypic and arkypallidal neurons, respectively. We found that opto-excitation of D2-SPN inputs with a light pulse of 1 ms reliably evoked inhibitory post-synaptic current (eIPSC) in nearly all prototypic neurons whereas a significant fraction of arkypallidal neurons (~30%) did not receive any inhibitory inputs from D2-SPNs (***Figure 1L***). In addition, the eIPSCs were on average 85% smaller in magnitude in arkypallidal than in prototypic neurons (***Figure 1K-L, supplementary table S1***). These properties were also confirmed when considering all putative GP neurons recorded (***Figure S4E-F, supplementary table S10***). Altogether, these results support the view that D2-SPN inputs differentially impact onto prototypic and arkypallidal neurons with arkypallidal neurons receiving weaker and less numerous projections from D2-SPNs as compared to prototypic neurons. Because prototypic neurons efficiently controlled BG network dynamics (Crompe et al., 2020) through their widespread local (i.e. intra GP) and distal projections (Bevan et al., 1998; Fujiyama et al., 2016; Kita and Kita, 1994; Mallet et al., 2012; Sato et al., 2000), their preferential inhibition by D2-SPNs will therefore cause a disinhibition in arkypallidal and STN neurons as revealed by their increased firing *in vivo*.

**Figure 1.**
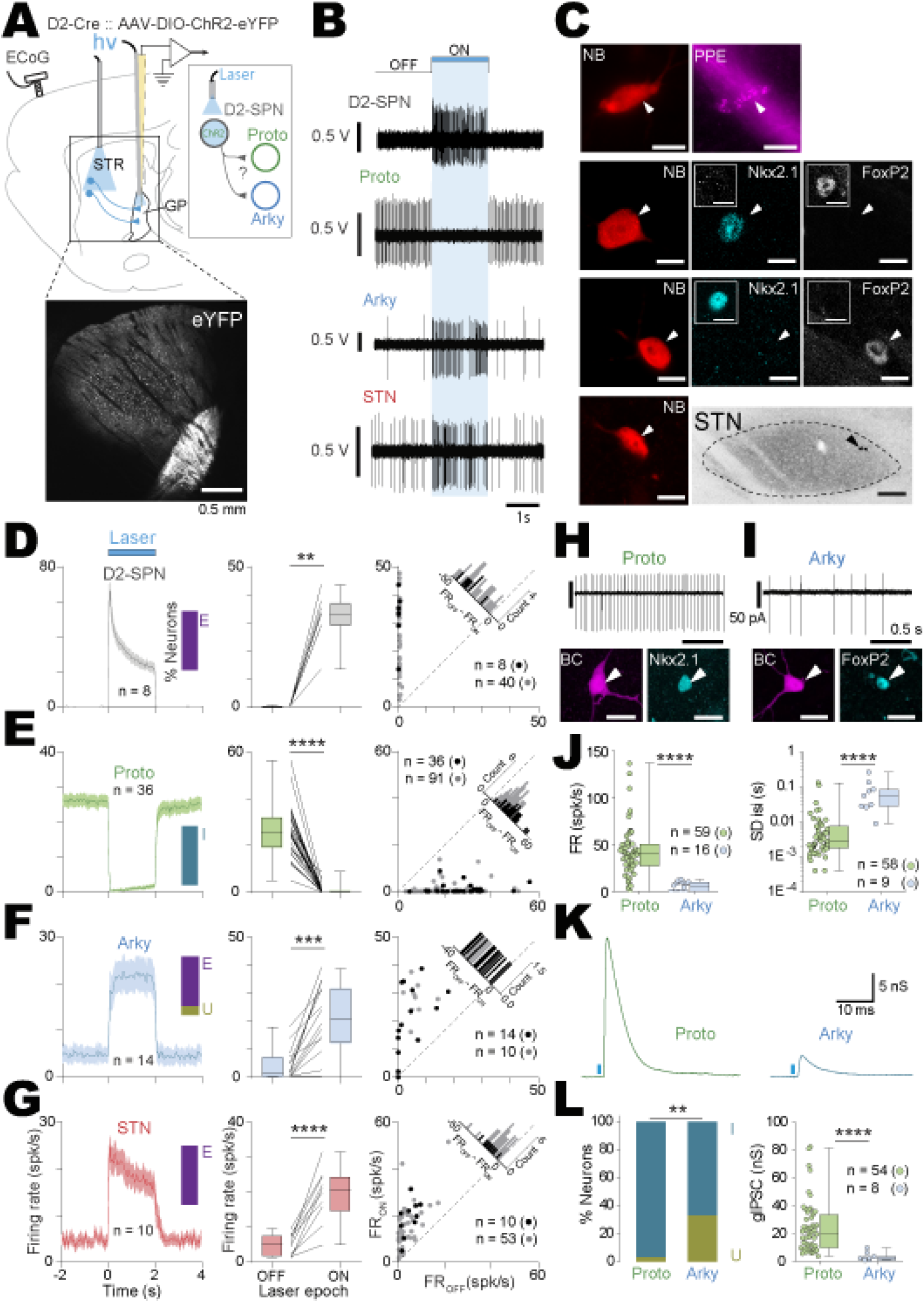
Opto-activation of D2-SPNs induced opposing effects on prototypic and arkypallidal neurons. ***(A)*** Schematic representation of the experimental design in D2-Cre mice injected with an AAV-DIO-ChR2-eYFP in the striatum (STR). The epifluorescent image illustrates the ChR2-eYFP expression in D2-SPNs with dense axonal projections to the GP (scale: 0.5 mm). ***(B)*** Representative examples of D2-SPN opto-excitation on the *in vivo* firing activity of identified D2-SPN, prototypic, arkypallidal and STN neurons. ***(C)*** Recorded neurons were juxtacellularly labelled with neurobiotin (NB) and then tested for the expression of specific molecular markers: D2-SPNs are preproenkephalin positive (PPE), prototypic and arkypallidal GP neurons express Nkx2.1 and FoxP2, respectively (scale bars confocal images: 10 μm, scale bar STN: 0.1mm). ***(D)*** Population PSTH (left, bin size: 50 ms), firing rate changes (middle), and scatter plot responses (right) during OFF vs. ON D2-SPN opto-excitation for all the recorded and juxtacellularly labelled (n= 8, black dots) or putative D2-SPNs (n = 40, grey dots). ***(E)*** Population PSTH (left, bin size: 50 ms), firing rate changes (middle), and scatter plot responses (right) during OFF vs. ON D2-SPN opto-excitation for all the recorded and juxtacellularly labelled (n= 36, black dots) or putative prototypic neurons (n = 91, grey dots). ***(F)*** Population PSTH (left, bin size: 50 ms), firing rate changes (middle), and scatter plot responses (right) during OFF vs. ON D2-SPN opto-excitation for all the recorded and juxtacellularly labelled (n= 14, black dots) or putative arkypallidal neurons (n = 10, grey dots). ***(G)*** Population PSTH (left, bin size: 50 ms), firing rate changes (middle), and scatter plot responses (right) during OFF vs. ON D2-SPN opto-excitation for all the recorded and juxtacellularly labelled (n= 10, black dots) or putative STN neurons (n = 53, grey dots). ***(H-I)*** Typical example of biocytin-filled prototypic (Nkx2.1+) and arkypallidal (FoxP2+) neurons with their corresponding activity recorded in cell-attached patch-clamp recordings (scale bars: 10 μm). ***(J)*** Box-and-whisker plots depicting average firing rate (FR) and the standard deviation of interspike interval (SDisi) of prototypic (green) and arkypallidal (blue) neurons. ***(K)*** Representative examples of inhibitory post-synaptic currents recorded in prototypic and arkypallidal neurons. ***(L)*** Bar graph representing the proportion of D2-SPN recipient prototypic and arkypallidal neurons and corresponding box-and-whisker plot of average IPSC conductance for each cell population. The inset bar plots in ***D-G*** represent the percentage of neurons excited (E), inhibited (I) and unaffected (U) by the light stimulation. Group data represents mean ± SEM, box-and-whisker plots indicate median, first and third quartile, min and max values. ** p<0.01, *** p<0.001, **** p<0.0001. See Table S1 for more details and statistical information.

### Opto-inhibition of STN neurons increased the firing rate of arkypallidal neurons *in vivo*

Although disinhibition mechanisms from prototypic neurons has been shown to account for the increased firing observed in the STN following GP opto-inhibition (Crompe et al., 2020; Kovaleski et al., 2020), the increased firing of arkypallidal neurons upon D2-SPN opto-excitation could alternatively be driven by the concurrent increase of STN firing. To better dissect the network mechanism and test if a disinhibition or a direct excitation is leading to an increased firing of arkypallidal neurons, we generated a D2-Cre∷ Vglut2-Cre double transgenic mouse line that provides the opportunity to opto-manipulate D2-SPNs and STN neurons independently in the same animal. We injected these mice with an AAV-DIO-ChR2 in the striatum—for opto-excitation of D2-SPNs—and an AAV-flex-ArchT into the STN—for opto-inhibition—(***Figure 2A***). We first validated the ArchT-mediated photo-inhibition of STN neurons (***Figure 2B***) and found that most STN neurons were indeed profoundly inhibited by ArchT activation (***Figure 2C, D and supplementary table S2***). We next tested the consequences of D2-SPN photo-excitation vs. STN photo-inhibition alone, or their concurrent effect on prototypic and arkypallidal neurons activity *in vivo* (***Figure 2E, F***). As expected from our previous experiments, opto-excitation of D2-SPN inputs induced opposite responses in the two GP cell populations with prototypic neurons showing an inhibition (***Figure 2G, I***) and with arkypallidal neurons increasing their activity (***Figure 2H, J***). Opto-inhibition of STN inputs also leads to opposite responses between the two GP cell-types with prototypic neurons decreasing their firing activity on average and arkypallidal neurons showing an overall increased response (***Figure 2I-J, supplementary table S2***). Interestingly, the suppression of STN tonic excitatory drive seems to affect the tonic firing of a small proportion of prototypic neurons whereas the arkypallidal response was a mirror image in the opposite direction. This highlights the putative contribution of collateral inhibition from prototypic neurons to shape the activity of arkypallidal neurons. The concurrent excitation of D2-SPN and inhibition of STN inputs did not suppress the excitatory response of arkypallidal neurons but only slightly decreased it (***Figure 2I-J, Supplementary Table S2**)*. Altogether, these results indicate that the response of arkypallidal neurons to D2-SPN inputs is principally driven by a disinhibition from prototypic neurons with only a small contribution from the direct STN excitatory drive.

**Figure 2.**
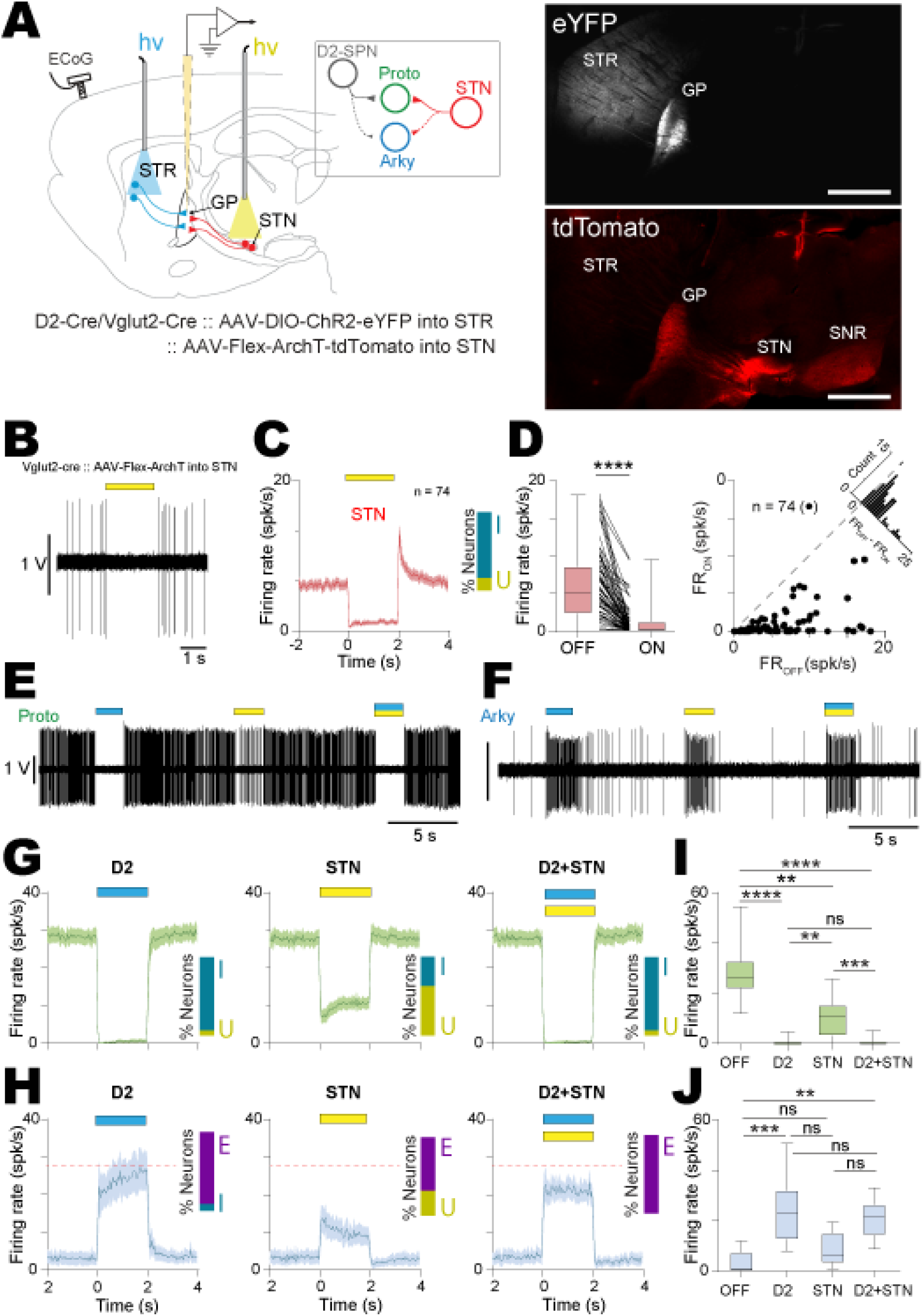
Opto-inhibition of STN neurons increased the firing rate of arkypallidal neurons *in vivo*. ***(A)*** Schematic representation of the experimental paradigm in double D2/Vglut2-Cre mice injected with an AAV-DIO-ChR2-eYFP in the striatum (STR) and an AAV-Flex-ArchT3-mCherry in the STN (scale bar: 2mm). ***(B)*** Representative opto-inhibition example of an STN neuron recorded *in vivo*. ***(C)*** Population PSTH (bin size: 50 ms) of STN neurons opto-inhibition (n = 74). ***(D)*** Box-and-whisker (left) and scatter plot distribution (right) of STN neurons firing during OFF *vs.* ON opto-inhibition (n = 74). ***(E, F)*** Representative firing activity examples of prototypic ***(E)*** or arkypallidal neurons ***(F)*** illustrating the effect of D2-SPN opto-activation (blue bar), STN opto-inhibition (yellow bar) and D2-SPN opto-activation + STN opto-inhibition (blue and yellow bar). ***(G, H)*** Population PSTHs (bin size: 50 ms) of D2-SPN opto-activation (left), STN opto-inhibition (centre) and D2-SPN opto-activation + STN opto-inhibition (right) effects on prototypic (n = 25) ***(G)*** and arkypallidal (n = 9) neurons ***(H)***. ***(I, J)*** Firing rate changes of prototypic (n = 25, ***I)*** or arkypallidal neurons (n=9, ***J)*** during OFF *vs*. ON D2-SPN opto-activation (D2), STN opto-inhibition (STN), and D2-SPN opto-activation + STN opto-inhibition (D2+STN). The inset bar plots in ***C, G,*** and ***H*** represent the percentage of neurons excited (E), inhibited (I) and unaffected (U) by the light stimulation. Group data represents mean ± SEM, box-and-whisker plots indicate median, first and third quartile, min and max values. ** p<0.01, *** p<0.001, **** p<0.0001, ns: not significant. See Table S2 for more details and statistical information.

### Subthalamic nucleus inputs preferentially excite prototypic neurons causing a disynaptic inhibition in arkypallidal neurons

To next define the functional connectome of STN inputs onto GP neurons, we used Vglut2-Cre mice injected with AAV-DIO-ChR2 in the STN (***Figure 3A***). Our *in vivo* electrophysiological recordings coupled with juxtacellular labeling confirmed the efficient opto-excitation of STN neurons in these mice, using either a 2 s (***Figure 3B-C***) or a 10 ms long laser pulses (***Figure S5A-B***). We then investigated the impact of STN inputs in GP *in vivo*. We found that all prototypic neurons were strongly excited by STN afferents (***Figure 3B, D-F, supplementary table S3)*** with fast latency (mean of 5.2 ± 0.6 ms) as best measured with the 10 ms laser stimulation protocols (***Figure S5A-D***). On the other hand, most arkypallidal neurons had a strong inhibition of their firing in response to STN inputs (***Figure 3B, D-F, supplementary table S3)*** which could last up to 50 ms when using short laser pulses (***Figure S5A-D, supplementary table S11***). These results were identical when considering a larger dataset that included all the putative GP neurons (***Figure S6, supplementary table 12***). Such differential firing responses toward STN inputs is consistent with our previous findings in rats in which opto-excitation of STN neurons led to quick excitation in prototypic neurons but inhibition in arkypallidal neurons (Crompe et al., 2020). Thus, similar to D2-SPN inputs, activation of STN glutamatergic afferents impact preferentially onto prototypic and not arkypallidal neurons. The strong excitatory response evoked in prototypic neurons is associated with an inhibition of arkypallidal neurons which was likely caused by disynaptic inhibition from prototypic axon collaterals. To investigate the synaptic origin of this *in vivo* differential GP neurons responses toward STN inputs, we next performed patch clamp recordings in acute brain slices of Vglut2-Cre mice injected with an AAV-DIO-ChR2 in the GP (***Figure 3G, H***). Interestingly, all identified prototypic and arkypallidal neurons that were recorded exhibited excitatory post-synaptic currents (EPSCs) evoked by opto-activation of STN synaptic terminals in the GP (***Figure 3I-J***). However, similar to inhibitory inputs coming from D2-SPNs, STN inputs were 74% smaller in arkypallidal than in prototypic neurons (***Figure 3J, supplementary table S3***). The same trend was also observed on a larger sample of unlabelled neurons (***Figure S7, supplementary table 13***). In conclusion, our data support the view that the functional connectome of STN inputs preferentially excites prototypic neurons that likely cause, in return, a powerful disynaptic inhibition in arkypallidal neurons. This result highlights the possibility that collateral inhibition from prototypic neurons is overriding a direct excitation of STN onto arkypallidal neurons and suggests that the activity of arkypallidal neurons is tightly controlled by prototypic neurons.

**Figure 3.**
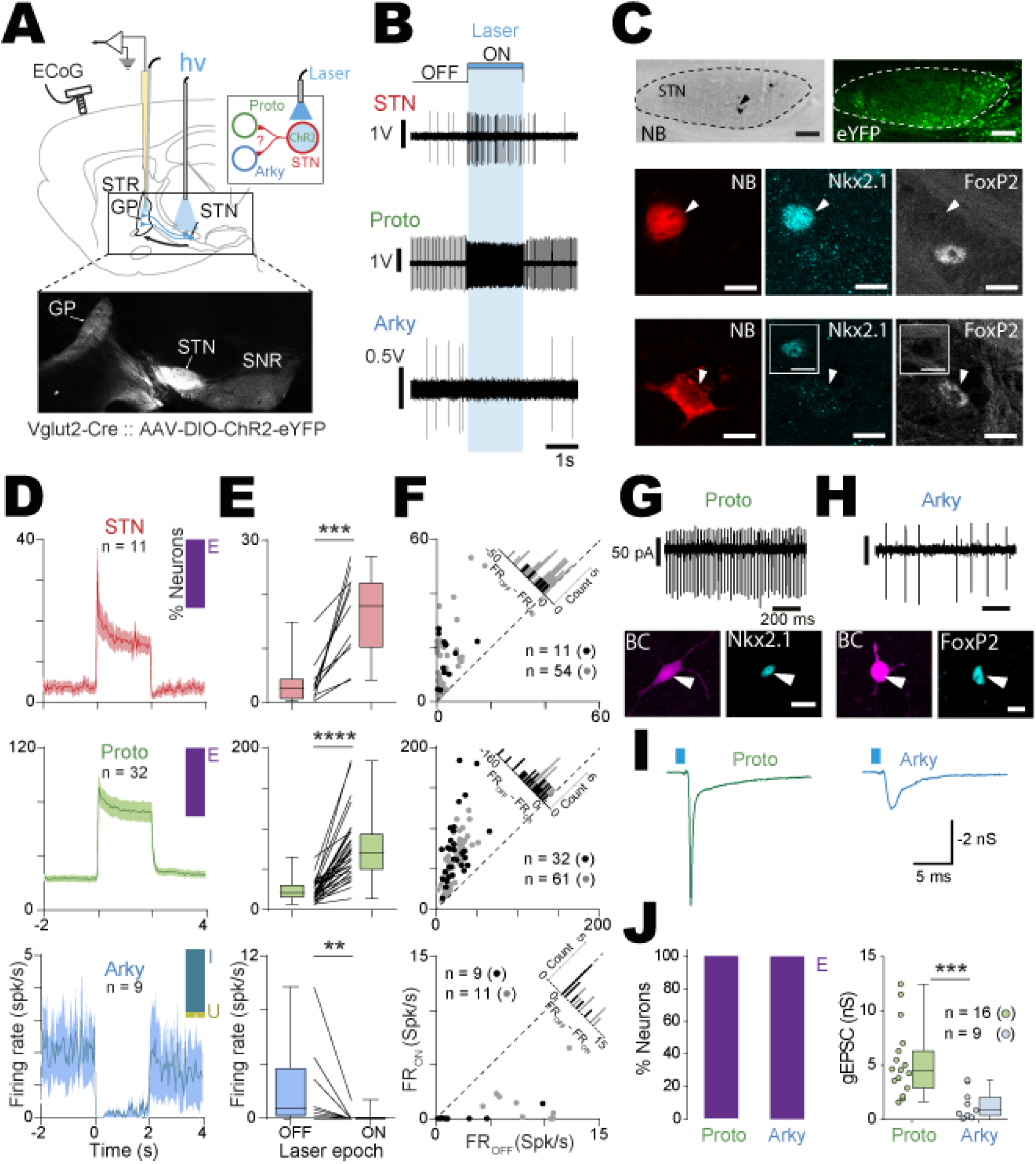
Opto-activation of STN neurons induced opposing effects on prototypic and arkypallidal neurons. ***(A)*** Schematic representation of the experimental design in Vglut2-Cre mice injected with an AAV-DIO-ChR2-eYFP in the STN, and recorded in the GP. The epifluorescent image is showing ChR2-eYFP expression in the STN and its axonal projections to the GP and SNR. ***(B)*** Representative examples of *in vivo* juxtacellularly labelled STN, prototypic and arkypallidal neurons showing the effect of STN opto-activation on their firing activities. ***(C)*** Microphotograph of STN (scale bars: 0.1 mm), prototypic and arkypallidal neurons juxtacellularly labelled *in vivo* with neurobiotin (NB, arrow) and tested for the labelling of Nkx2.1 and FoxP2 (scale bar confocal images: 10 μm). ***(D)*** Population PSTH (bin size: 50 ms) of all the *in vivo* juxtacellularly labelled STN (n = 11, top), prototypic (n = 32, middle), and arkypallidal (n = 9, bottom) during STN opto-activation. The inset bar plots represent the percentage of neurons excited (E), inhibited (I) and unaffected (U) by the light stimulation***. (E)*** Box-and-whisker plots of the firing rate changes of all the juxtacellularly labelled STN (n = 11, top), prototypic (n = 32, middle), and arkypallidal (n = 9, bottom) during laser OFF *vs.* ON STN opto-activation. ***(F)*** Scatter plot distributions of the juxtacellularly labelled and putative STN (top, n = 11, black dots and n = 54, grey dots, respectively), prototypic (middle, n = 32, black dots and n = 61, grey dots, respectively), and arkypallidal neurons (bottom, n = 9, black dots and n = 11, grey dots, respectively). (***G, H)*** Biocytin-filled prototypic (Nkx2.1+) and arkypallidal (FoxP2+) neurons with their corresponding activity recorded in cell-attached patch-clamp recordings. ***(I)*** Representative examples of excitatory post-synaptic currents recorded in prototypic and arkypallidal neurons. ***(J)*** Bar graphs representing the proportion of STN recipient prototypic and arkypallidal neurons and corresponding box-and-whisker plot of average EPSC conductance for each cell population. Group data represents mean ± SEM, box-and-whisker plots indicate median, first and third quartile, min and max values. ** p<0.01, *** p<0.001, **** p<0.0001. See Table S3 for more details and statistical information.

With this in mind, we next tested if the disynaptic inhibition of prototypic neurons onto arkypallidal neurons could be suppressed by external inputs, thus unmasking the direct excitation of STN inputs onto arkypallidal neurons. To do so, we studied the effect of concurrent opto-excitation of D2-SPN inputs (to suppress the activity of prototypic neurons) and STN inputs (to drive excitation) on the activity of arkypallidal neurons in D2-Cre∷ Vglut2-Cre mice injected with the AAV-DIO-ChR2 virus in both the striatum and STN (***Figure S8A, B***). As described before, photo-activation of D2-SPN inputs increased the firing of arkypallidal neurons (***Figure S8C***), while photo-activation of the STN induced an overall inhibition of their activity (***Figure S8D)*** and these results were similar for both long (i.e. 2s) and short (i.e. 10 ms) laser stimulation protocols. Interestingly though, the concurrent opto-activation of both D2-SPN and STN inputs induced a stronger excitation in arkypallidal neurons than D2-SPN inputs alone (***Figure S8E***), although this effect was not statistically significant in our sample size (***Figure S8F-G, supplementary table S14***). We have previously shown that the excitatory response of arkypallidal neurons induced by D2-SPN inputs is mostly caused by a disinhibition mechanism with only a small contribution of the STN direct excitation (see ***Figure 2***). We thus confirm here that STN inputs are capable of directly exciting arkypallidal neurons but only when the disynaptic inhibition normally exerted by prototypic neurons is removed. This suggests a rather complex mechanism by which striatal D2-SPN inputs could gate the integration of STN information in the GP and determine which cell population in the GP will be excited by STN inputs.

### Prototypic neurons control the activity of arkypallidal neurons through axon collateral inhibition

GP neurons, especially prototypic neurons, have dense axon collaterals (Bugaysen et al., 2013; Mallet et al., 2012; Miguelez et al., 2012; Sadek et al., 2007; Sato et al., 2000), yet characterization of lateral inhibition between prototypic and arkypallidal neurons has never been investigated functionally. To first explore if prototypic neuron collaterals could efficiently inhibit arkypallidal neurons, we injected an AAV-DIO-ChR2 virus into the GP of Nkx2.1-Cre mice that provides genetic access to the prototypic population based on the selective expression of Nkx2.1 in prototypic neurons (Abdi et al., 2015; Dodson et al., 2015). The expression of ChR2 as reported by the visualization of the reported protein mCherry, was accordingly restricted to prototypic neurons (***Figure 4A-B***). Opto-excitation of prototypic neurons in cell-attached configuration, produced a strong and sustained increase in their firing (***Figure 4C, E, supplementary table S4***), while it induced a potent inhibition of the firing activity of all recorded arkypallidal neurons (***Figure 4D, F, supplementary table S4***). Light-evoked IPSCs were detected both in arkypallidal and prototypic neurons. Also our sample size did not yield to a significant difference, our data trend towards larger IPSCs in arkypallidal neurons (***Figure 4G, supplementary table S4***). These results support the existence of functional collateral connections between prototypic and arkypallidal neurons, as previously suggesting by anatomical studies (Mallet et al., 2012). We then investigated if these collateral connexions were reciprocal with arkypallidal inhibiting prototypic neurons. To do so, we injected an AAV-DIO-ChR2 into the GP of FoxP2-Cre mice (***Figure 5A***) that enabled specific expression of ChR2 in arkypallidal neurons (***Figure 5B***) and recorded GP neurons in acute brain slices. Photo-activation of arkypallidal neurons efficiently increases the firing in all recorded arkypallidal neurons thus validating this approach (***Figure 5C, E, supplementary table S5***), however, this strategy did not produce any inhibitory responses in prototypic neurons (***Figure 5D, F, supplementary table S5***). To confirm that the absence of inhibitory responses was genuinely caused by a lack of connections rather than an artefactual disconnection produced by the slicing procedure, we next performed *in vivo* single unit recordings in anaesthetized FoxP2-Cre mice similarly injected with an AAV-DIO-ChR2 in the GP (***Figure 5G***). Light-activation also led to an increased firing rate of arkypallidal neurons recorded *in vivo (**Figure 5H**)*, but this excitation had no effect on the average output activity of GP prototypic neurons (***Figure 5I, supplementary table S5)***. Looking further into the effects at individual neuron level, we found that photo-excitation of arkypallidal neurons produces for the most part no effect on prototypic neurons whereas a minority was either excited or inhibited (***Figure 5I***), supporting the conclusion of a rather weak functional connection between arkypallidal and prototypic neurons. Altogether, our results are in line with the small proportion of arkypallidal neurons (~30%) compared to the total number of GP neurons (Dodson et al., 2015) and the small number of synaptic terminals made by these neurons within the GP (Mallet et al., 2012). This dataset thus suggests the existence of a unilateral connectivity from prototypic to arkypallidal neurons which provides the anatomical architecture to promote a disynaptic circuit capable of turning the activity of arkypallidal neuron ON or OFF.

**Figure 4.**
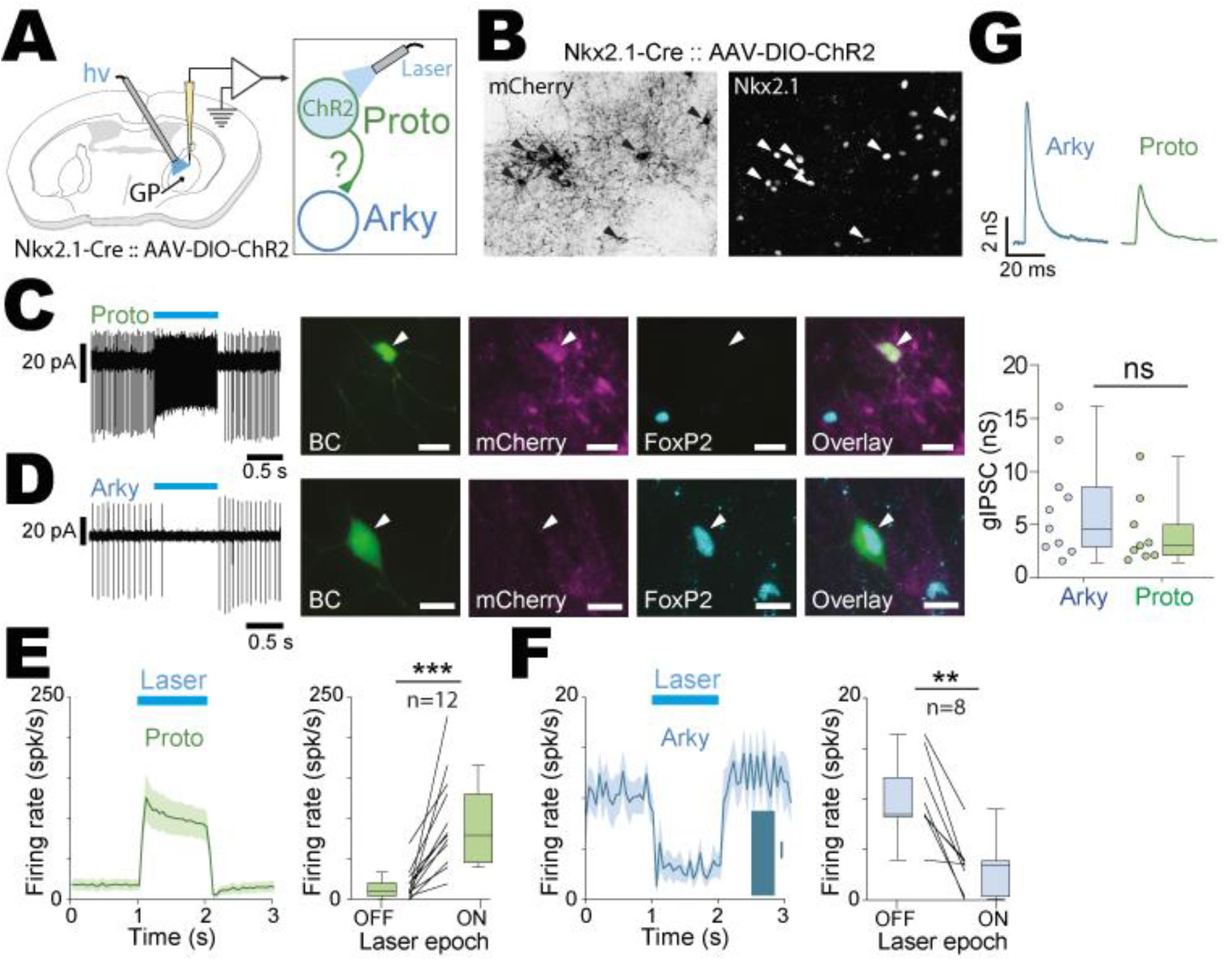
Prototypic neuron axon collaterals powerfully inhibit arkypallidal neurons. ***(A)*** Schematic representation of the experimental design for the *ex vivo* experiments. ***(B)*** Histological verification of the expression of the AAV5-EF1a-DIO-ChR2(H134R)-mCherry in prototypic neurons (Nkx2.1+). ***(C)*** Typical cell-attached activity of an identified prototypic neuron during OFF and ON epochs of laser stimulation with corresponding confocal images of a prototypic neuron filled with biocytin (BC) and immuno-positive for Nkx2.1 (scale bars:10 μm). ***(D)*** Typical cell-attached activity of an identified arkypallidal neuron inhibited during ON epoch of laser stimulation with corresponding confocal images of an arkypallidal neuron filled with biocytin (BC) and immuno-positive for FoxP2 (scale bars:10 μm). ***(E)*** Population PSTH of prototypic neurons firing during opto-activation (bin: 50 ms). Box-and-whisker plots representing average firing rate during ON-OFF epochs of laser stimulation. Each line indicates a single neuron change in firing rate. ***(F)*** Population PSTH of arkypallidal neurons firing inhibition during opto-activation of prototypic neurons (bin: 50 ms). Box-and-whisker plots representing the average firing rate during ON-OFF epochs of laser stimulation. Each line indicates a single neuron change in firing rate. ***(G)*** Representative examples of GABAergic IPSC recorded in arkypallidal (Blue) and prototypic (Green) neurons. Box-and-whisker plots representing IPSC conductances in each cell population. Group data represents mean ± SEM, box-and-whisker plots indicate median, first and third quartile, min and max values. ** p<0.01, ns: not significant. See Table S4 for more details and statistical information.

**Figure 5.**
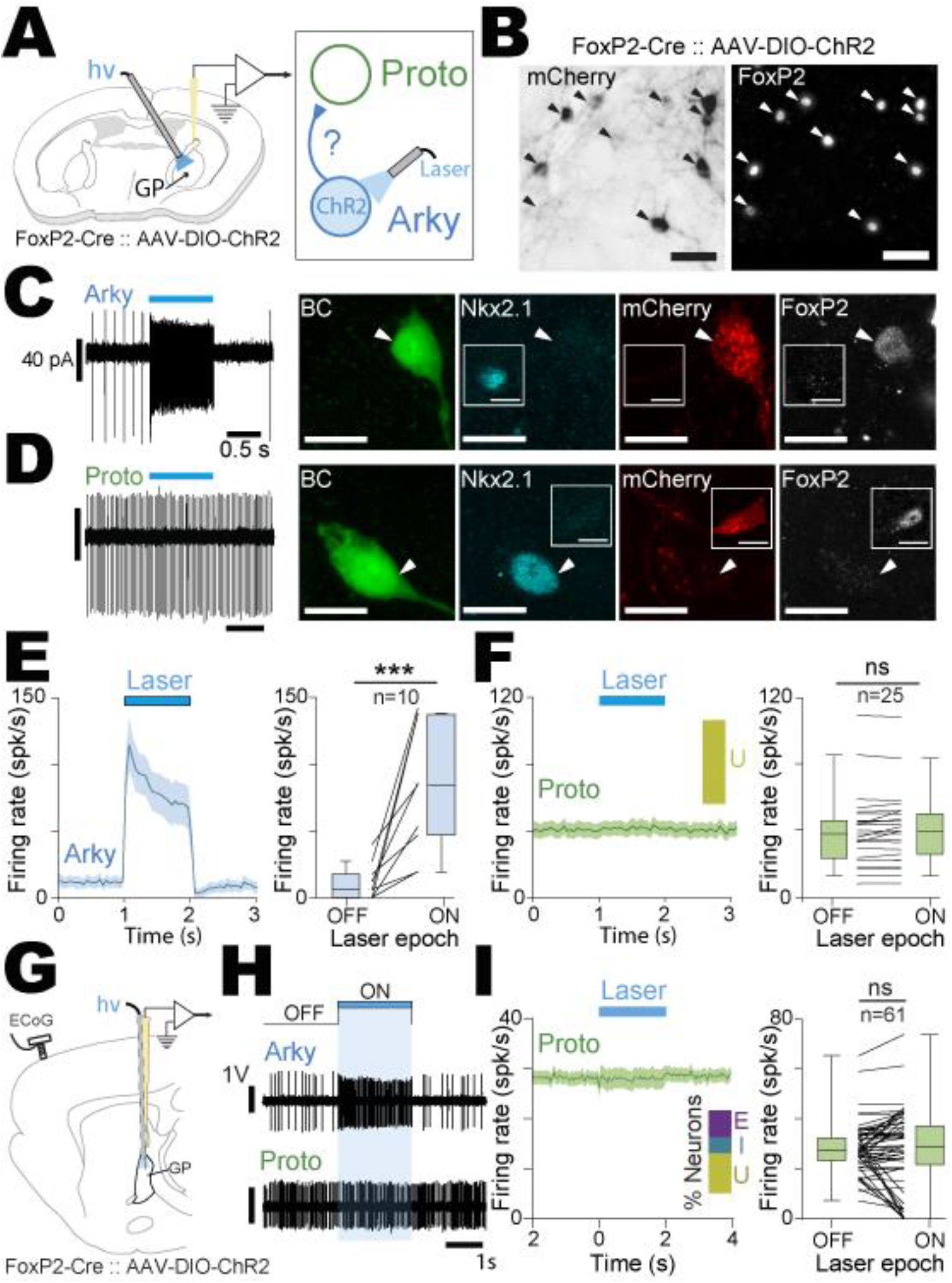
Arkypallidal neurons do not provide functional inhibition onto prototypic GP neurons. ***(A)*** Schematic representation of the experimental design for the *ex vivo* experiments. ***(B)*** Histological verification of the expression of the AAV-DIO-ChR2-mCherry in arkypallidal neurons (FoxP2+). ***(C)*** Typical cell-attached activity of an identified arkypallidal neuron during OFF and ON epochs of laser stimulation with corresponding confocal images of the biocytin-filled (BC) neuron and immuno-positive for FoxP2. ***(D)*** Typical cell-attached activity of an identified prototypic neuron which activity remains unchanged during the ON epoch of laser stimulation with corresponding confocal images of a prototypic neuron filled with biocytin (BC) and immuno-positive for Nkx2.1. ***(E)*** Population PSTH of arkypallidal neurons firing during opto-activation (bin: 50 ms). Box-and-whisker plots representing the average firing rate during ON-OFF epochs of laser stimulation. ***(F)*** Population PSTH (left, bin size: 50 ms) of prototypic neurons firing during opto-activation of arkypallidal neurons. Box-and-whisker plots (right) representing the average firing rate during ON-OFF epochs of laser stimulation. ***(G)*** Schematic representation of the experimental design for the *in vivo* experiments. ***(H)*** Typical examples of the activity of an arkypallidal neuron and a prototypic neuron during ON/OFF epochs of light stimulation. ***(I)*** Population PSTH (left, bin size: 50 ms) of firing rate changes of all the recorded prototypic neurons during opto-activation of arkypallidal neurons. Box-and-whisker plots (right) representing the average firing rate of prototypic neurons during ON-OFF epochs of laser stimulation. The inset bar plots in ***F*** and ***I*** represent the percentage of prototypic neurons excited (E), inhibited (I) and unaffected (U) by the light stimulation. Scale bars represent 50 μm (in ***B)***, or 10 μm (in ***C***, ***D***). Group data represents mean ± SEM, box-and-whisker plots indicate median, first and third quartile, min and max values. ** p<0.01, ns: not significant. See Table S5 for more details and statistical information.

### Excitation of arkypallidal neurons is sufficient for locomotion inhibition

Activation of both D2-SPNs (i.e. the indirect ‘No Go’ pathway) and STN neurons (i.e. the hyperdirect ‘Stop’ pathway) has been considered to favor motor suppression. Considering the opposite impact that both pathway stimulations had on GP prototypic and arkypallidal neuronal firing, we next tested if the motor correlates caused by the photoactivation of both circuits were identical. To best detect an effect on movement inhibition, we recorded the locomotor activity of mice walking on a motorized treadmill and applied random 500 ms duration optogenetic stimulations (***Figure 6A***). Both indirect (i.e. D2-SPNs) and hyperdirect (i.e. STN) pathways were photo-activated using bilateral injections of an AAV-DIO-ChR2 virus in the striatum (***Figure 6B***) or the STN (***Figure 6C***) of D2-Cre and Vglut2-Cre mice, respectively. Interestingly, we found that D2-SPN and STN stimulations caused different effects on locomotion. Indeed, activation of D2-SPNs induced a pronounced reduction in movement velocity ***(Figure 6E, I, supplementary table 6)*** whereas the STN stimulation had no clear effect ***(Figure 6F, I, supplementary table 6***). Locomotion experiments using mice injected with a control AAV-DIO-eYFP virus were also performed to test for the potential side effects induced by the light and the virus transduction associated with our optogenetic manipulations. These control optogenetic experiments show no effect of light stimulation on the locomotor behaviour (***Figure 6H, I, supplementary table 6***). We next dissected whether the differential neuronal response observed in prototypic and arkypallidal neurons upon D2-SPN or STN stimulations could account for the differential effect on locomotion. Indeed, D2-SPN stimulation induced a pause in prototypic neurons firing and a strong disynaptic excitation in the arkypallidal population. Because arkypallidal neurons have been proposed to play an important role in movement cancellation (Mallet et al., 2016), we hypothesized that the locomotion inhibition consequent to D2-SPN stimulations was mediated by the increased firing of arkypallidal neurons. To tease apart the specific functional contribution of prototypic vs. arkypallidal neuronal response in locomotion inhibition, we used FoxP2-Cre mice as these transgenic mice gave us the opportunity to test the contribution of arkypallidal excitatory response without affecting the overall firing of prototypic neurons (see results of ***Figure 5***). Interestingly, we found that optogenetic excitation of arkypallidal FoxP2-expressing neurons reproduced a strong inhibition of locomotion ***(Figure 6D, G, I, supplementary table 6)*** and that this effect was significantly different in all FoxP2-Cre mice as compared to matched controlled experiments. Altogether, our results suggest that increased activity in arkypallidal neurons is sufficient to mediate locomotion inhibition.

**Figure 6.**
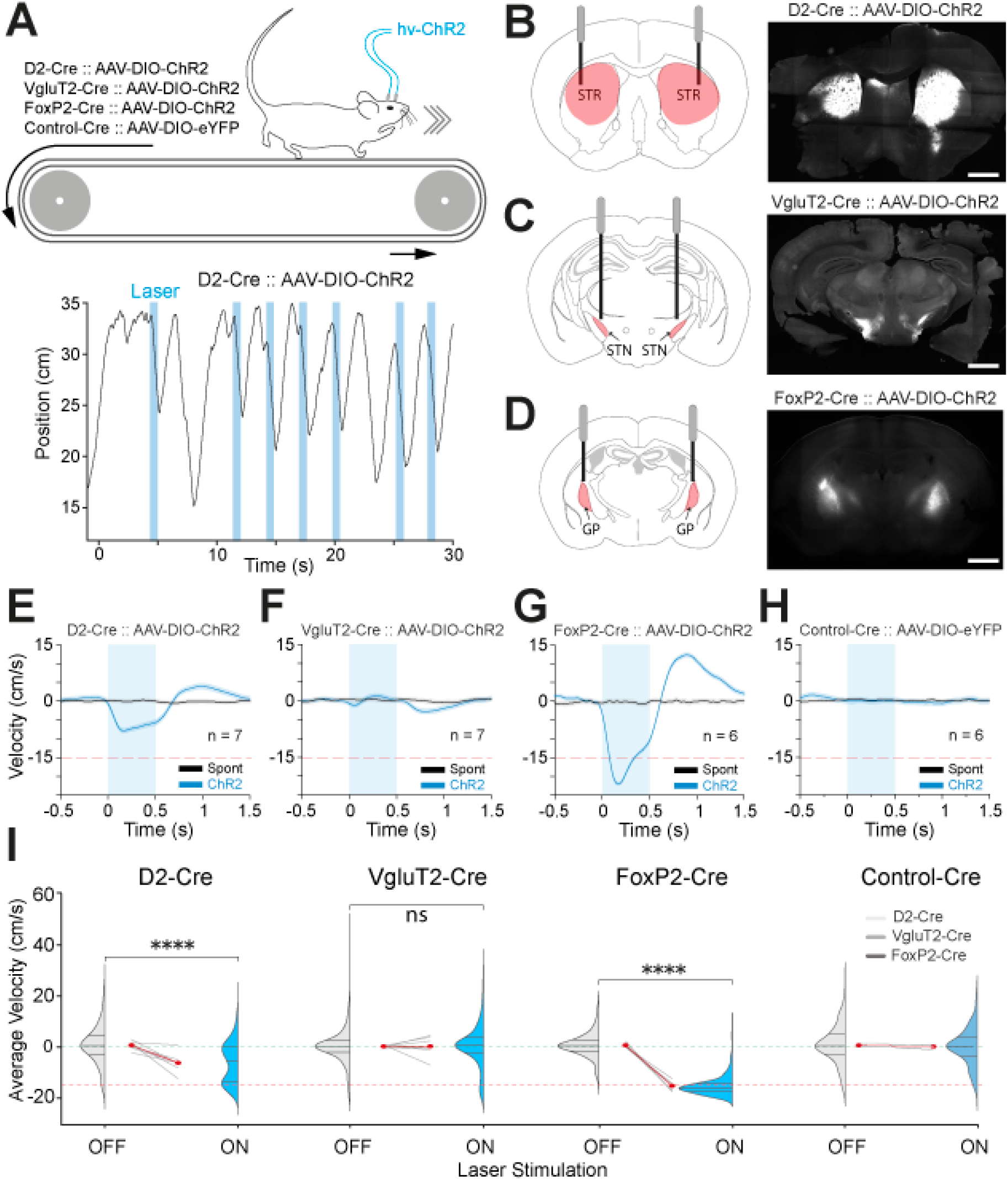
Activation of arkypallidal neurons is sufficient to induce locomotion inhibition. ***(A)*** Schematic representation of the experimental paradigm for behavioural analysis of the locomotion in mice running on a motorized treadmill. D2-Cre, Vglut2-Cre, FoxP2-Cre mice were injected bilaterally with an AAV-DIO-ChR2-eYFP or a control AAV-DIO-eYFP in the striatum (n=7 and n=2, respectively), the STN (n=7 and n=2, respectively), and in the GP (n=7 and n=2, respectively). Raw example (bottom) of 30 s locomotion bouts in a D2-Cre∷AAV-ChR2 mice receiving blue light stimulation (500 ms long) at random timing. ***(B-D)*** Brain schematics with optic fibre positions in the target areas and corresponding epifluorescent images illustrating the histological verification of the ChR2-eYFP virus expression in D2-Cre ***(B)***, Vglut2-Cre ***(C)***, and FoxP2-Cre ***(D)*** mice (scale bars represent 1 mm) ***(E-F)*** Average velocity induced by the light stimulation (blue) or during spontaneous locomotion (black) in D2-Cre∷AAV-ChR2 ***(E)***, Vglut2-Cre∷AAV-ChR2 ***(F)***, FoxP2-Cre∷AAV-ChR2 ***(G)***, and all the various control mice experiments (Control-Cre∷AAV-eYFP, ***H***). ***(I)*** Velocity population graphs showing the velocity distributions during laser ON/OFF in the different animals. The grey lines represent individual animals and the red lines represent the mean between animals. The dotted red line shows the velocity of the treadmill (15 cm/s). Group data represents mean ± SEM, distribution plots indicate median, first and third quartile, min and max values. **** p<0.0001, ns: not significant. See Table S6 for more details and statistical information.

## Discussion

### Increased arkypallidal activity causes locomotion inhibition

It has been proposed that arkypallidal neurons broadcast a “stop” signal (***Figure S9A)*** to the striatum when an ongoing motor action has to be cancelled (Mallet et al., 2016). However, the direct contribution of arkypallidal neurons in motor inhibition has never been interrogated with optogenetic stimulations before. Here, we show that opto-excitation of arkypallidal neurons causes a robust inhibition of on-going locomotor activity. Interestingly, the light-driven activity increase of arkypallidal neurons does not modify the overall output activity of prototypic neurons. This suggests that the arkypallidal-mediated inhibitory effect on movement is independent of any changes in prototypic neurons activity and thus, their downstream projection sites. Instead, we believe that locomotion inhibition is caused through a global suppression of ‘Go-related’ striatal activity mediated by the dense projections sent from arkypallidal neurons to the striatum (Fujiyama et al., 2016; Mallet et al., 2012) and which contact both striatal interneurons and SPN cell populations (Mallet et al., 2012). These results reveal an important function of arkypallidal neurons that is to control movement-related activity in the striatum either to suppress on-going movement or to inhibit competitive actions. It is possible that action initiation is promoted by the inhibition of arkypallidal neurons. However, recordings of arkypallidal neurons during spontaneous movements have mainly shown increased rather than decreased activity in this cell population (Dodson et al., 2015). That being said, it is possible that their low firing activity in awake animals makes inhibition less detectable in arkypallidal neurons than in fast firing neurons. Interestingly, it has been described that D1-SPNs send bridging collaterals to the GP and that structural plasticity in these connections can rescue locomotor imbalance (Cazorla et al., 2014). If the function of D1-SPNs in movement execution is principally driven by their direct inhibitory effect onto SNR activity (Kravitz et al., 2010; Roseberry et al., 2016), it is also possible that the bridging collaterals of D1-SPNs specifically suppress the activity of arkypallidal neurons. Thus, inhibiting the motor brake function exerted by arkypallidal neurons might facilitate action initiation and motor sequence execution. In support of this hypothesis, a recent pre-printed work shows that the D1-SPN inhibitory inputs have a strong bias towards arkypallidal neurons (Ketzef and Silberberg, 2020).

### D2-SPNs opto-activation leads to a disynaptic excitation of arkypallidal neurons

In this study, we found that D2-SPN optogenetic activation induces locomotion inhibition in mice walking on a motorized treadmill. This result confirms the negative influence of D2-SPN activity on movement execution (Cui et al., 2013; Durieux et al., 2009; Kravitz et al., 2010; Oldenburg and Sabatini, 2015; Panigrahi et al., 2015; Sheng et al., 2019; Tecuapetla et al., 2016). Striatal D2-SPNs are one of the main GABAergic inputs to the GP (Kita, 2007) that inhibit GP neurons (Miguelez et al., 2012; Sims et al., 2008). Our *ex vivo* electrophysiological recordings demonstrate that both prototypic and arkypallidal neurons receive inhibitory inputs from D2-SPNs, although the inhibitory currents detected in arkypallidal neurons are smaller than the ones in prototypic neurons. Such differential striatal innervation onto GP neurons has already been reported but the molecular identity of GP neurons was not defined (Chuhma et al., 2011; Kita and Kitai, 1991). By further dissecting how D2-SPN inputs impact onto different GP neuronal populations *in vivo*, we revealed a cell-type specific integration of these inputs. Indeed, prototypic neurons were almost fully silenced by D2-SPN inputs, whereas arkypallidal neurons were strongly excited. This opposition of response in arkypallidal neurons was caused by a disynaptic disinhibition mechanism involving axon collaterals from prototypic neurons. These results outline how a difference in the organization of synaptic inputs at the single-cell level measured *ex vivo* can give rise to opposite firing activities when cell populations are embedded into neuronal networks *in vivo*. In addition, our finding regarding D2-SPN stimulation leading to an increase in the activity of arkypallidal neurons raises new insights on the cellular mechanism causing movement inhibition consequent to the indirect pathway activation. In particular, we show that the excitation of arkypallidal neurons induces locomotion inhibition and this effect is certainly due to the global suppression of striatal activity (***Figure S9B***). Although we show in this work that this mechanism is sufficient by itself to suppress locomotion, it is possibly acting in concert with the inhibition of prototypic neurons that rather promote movement inhibition through an increased activity of BG outputs. With this in mind, movement inhibition induced by the stimulation of the indirect pathway might thus be relying on this dual but synergistic disynaptic mechanism: the classic increased activity of BG output (i.e. through the suppression of prototypic neurons firing) and, in parallel, a suppression of ‘Go-related’ striatal activity (i.e. through the increased firing of arkypallidal neurons). Altogether, the present findings highlight new concepts on the motor function of the indirect pathway. Whether this disynaptic circuit mechanism is also used for basal ganglia non-motor function is not known but it is tempting to speculate that it represents a key mechanism by which D2-SPNs can modulate D1-SPNs activity (Matamales et al., 2020).

### STN inputs cause a disynaptic inhibition in arkypallidal neurons and can be gated by D2-SPN inputs

The STN is the only glutamatergic nucleus in the BG, and it represents the main excitatory input to GP neurons (Kita and Kitaï, 1987; Smith et al., 1990). It has been shown in mice that optogenetic activation of STN axon terminals within the GP increases glutamate release (Viereckel et al., 2018) and c-Fos expression (Tian et al., 2018). We show here that, similar to D2-SPN inputs, STN inputs differentially impact the activity of prototypic and arkypallidal neurons. Our *ex vivo* data demonstrate that both cell populations receive STN excitation but the excitatory currents are smaller in arkypallidal neurons as compared to prototypic neurons. This differential input organization translates into opposite firing responses when studied *in vivo*: STN opto-excitation induces an increased activity of prototypic neurons and a decreased firing in arkypallidal neurons. This opposition in firing rates is in agreement with previous findings in rats (Crompe et al., 2020). However, we further elucidate here the cellular mechanisms responsible for this opposition of response. Indeed, STN inputs preferentially excite prototypic neurons that, in turn, inhibit arkypallidal neurons through their powerful axon collaterals. This disynaptic mechanism is also revealed when testing the consequences of STN opto-inhibition. More specifically, we found that a proportion of prototypic neurons responds to STN inhibition with a partial reduction in their firing activities which is sufficient to cause a disinhibition in arkypallidal neurons. Noteworthy, this excitatory effect was weaker than the one induced by D2-SPN stimulation. The rather limited effect of STN inhibition on prototypic neuronal activity could be explained by the autonomous pacemaking firing properties of prototypic neurons (Mallet et al., 2019), and is consistent with previous work (Kovaleski et al., 2020). Altogether, these results confirm the powerful inhibitory control exerted by prototypic neurons axon collaterals onto arkypallidal neurons upon STN opto-excitation. Interestingly, we did not find any clear effect of STN opto-excitation when looking at the motor performance of mice walking on a treadmill. This result might appear surprising at first but is consistent with previous findings supporting that STN opto-stimulation produces hyperkinetic behaviors similar to the ones induced by the direct excitation of GP neurons (Tian et al., 2018). It is thus likely that STN excitation competes with the STN-induced prototypic increased activity at the level of BG outputs to control movement execution. Moreover, the disynaptic inhibition induced in arkypallidal neurons upon STN opto-excitation will leave striatal ‘Go-related’ signals without arkypallidal inhibitory control (***Figure S9C)***. In addition, our experimental data obtained when co-exciting D2-SPN and STN inputs, suggest that D2-SPN neurons are gating the integration of STN inputs in arkypallidal neurons. Indeed, we found that the simultaneous activation of D2-SPNs and STN neurons produces a switch from inhibition to robust excitation in the firing of arkypallidal neurons that is stronger than D2-SPNs opto-excitation alone. The end result of this co-activation suggests that STN neurons can now influence the firing of arkypallidal neurons and thus provide a ‘stop signal’ mediating behavioral inhibition. Importantly, the dysfunction of this gating mechanism could be the source of the excessive inhibition of movements present in parkinsonism and that has been attributed to the hyperactivity of STN neurons (Bergman et al., 1990).

### Prototypic GP axon collaterals act as a switch for arkypallidal neuron activity

Previous work have shown that GP neurons send axon collaterals to other GP neurons (Kita and Kita, 1994; Sadek et al., 2007; Sato et al., 2000) and that prototypic neurons project axon collaterals onto arkypallidal neurons (Mallet et al., 2012). Here, we demonstrate that the axon collaterals from prototypic neuron powerfully control arkypallidal neuron activity. Interestingly, the collaterals connections from arkypallidal to prototypic neurons are not reciprocal or only targets a subset of prototypic neurons that were not sampled in our *ex vivo* recordings. Indeed, the population of prototypic neurons has been further divided into subclasses based on the differential expression of molecular markers (Abdi et al., 2015; Abecassis et al., 2020; Mastro et al., 2014, 2017). Another important aspect for future work is to determine if specific inputs can directly control the activity of arkypallidal neurons (Abecassis et al., 2020; Hunt et al., 2018; Karube et al., 2019) or whether their activity is always disynaptically controlled by prototypic axon collaterals. Altogether, our work reveals a novel and unaccounted for disynaptic motif circuit in the GP that represents an efficient mechanism by which indirect pathway stimulation could inhibit ongoing actions. Our results thus call for a re-evaluation of the BG functional organization that should account for the key role played by arkypallidal neurons to control striatal circuits for action selection and cancellation.

## Methods

### Animals

All Experimental procedures were performed on both male and female adult (>6 weeks) mice with C57BL/6J genetic backgrounds in accordance with the European legislation for the protection of animals used for scientific purposes (directive 2010/63/EU). D2-SPNs were specifically targeted using D2-Cre mouse line (GENSAT, catalog no. 032108-UCD), STN neurons were targeted using Vglut2-Cre mouse line (The Jackson Laboratory, stock no. 016963), prototypic GP neurons were targeted using Nkx2.1-Cre mouse line (The Jackson Laboratory, stock no. 008661), and arkypallidal GP neurons were targeted using the FoxP2-Cre mouse line (The Jackson Laboratory, stock no. 030541). To enable targeting of both D2-SPNs and STN neurons in the same animals, we generated a D2-Cre∷ Vglut2-Cre mouse line by breeding hemizygous Vglut2-Cre driver mice with homozygous D2-Cre mice. To promote ChR2 expression in all D2-SPNs we generated a D2-Cre∷ Ai32 mice by breeding D2-cre driver mice with homozygous Ai32 mice (The Jackson Laboratory, stock no. 024109). This project was approved by the French ministry of higher education and research and the ethical committee of CNRS, Aquitaine Region (accreditation numbers #6231 and #25001). Mice were housed collectively (at least by 2 per cage) under artificial conditions of light (light/dark cycle, light on at 7:00 a.m.), temperature (24°C), and humidity (45%) with food and water available ad libitum. A total number of 151 mice were used for this study of which 12 mice were excluded due to failed or missed virus injection associated both with low transduction rate of targeted area.

### Stereotaxic injection of viral vectors

All the virus used for our optogenetic manipulations were adeno-associated virus directly purchased from a vector core (UNC vector core) and micro-injected under stereotaxic condition in D2-Cre, Vglut2-Cre, double D2-Cre∷ Vglut2-Cre, Nkx2.1-Cre, and FoxP2-Cre mice using glass capillary (tip diameter 35 μm; 1-5 μL Hirschmann^®^ microcapillary pipette, CAT#Z611239, Sigma-Aldrich) connected to a Picospritzer^®^ pressure system (Parker Hannifin). Briefly, mice were anesthetized with isoflurane (induction/maintenance: 3-5/1.5%; Iso-vet^®^, Piramal healthcare), fixed on a stereotaxic frame (Unimécanique, M2e) and placed on heating blanket. An ophthalmic ointment (Liposic^®^, Bauch & Lomb Swiss) was used all along the surgery to prevent dehydration. Photo-stimulation of D2-SPNs or STN neurons were performed using an AAV5-EF1α-DIO-ChR2(H134R)-eYFP viral vector (4×10^12^ viral particles/mL) injected in D2-Cre mice (n = 80, 2 points of injections in dorsolateral striatum at coordinates AP: 0.8 mm and 0.3 mm anterior to bregma, ML: 1.75 mm and 2.2 mm, DV: 2.6 mm, volume: 400 nL per sites) or Vglut2-Cre mice (n = 61, injection in STN at coordinates AP: 1.8 mm posterior to bregma, ML: 1.5 mm, and DV: 4.5 mm, volume: 150 nL per sites). Photo-inhibition of STN neurons were performed using an AAV5-CAG-Flex-ArchT-tdTomato (6×10^12^ viral particles/mL), injected in the STN of Vglut2-Cre mice (n = 8, same coordinates as above). Combined injections of these viruses were used for dual photo-excitation of both D2-SPNs and STN inputs (n = 12 D2-Cre∷ Vglut2-Cre mice) or photo-excitation and photo-inhibition of D2-SPNs and STN inputs, respectively (n = 10 D2-Cre∷ Vglut2-Cre mice). Photo-excitation of prototypic or arkypallidal neurons were performed using an AAV5-EF1α-DIO-ChR2(H134R)-mCherry (5.2×10^12^ viral particles/mL) injected in the GP (coordinates AP: 0.35 mm posterior to bregma, ML: 2.0 mm, DV: 3.6 mm, volume: 120-200 nL per sites) of Nkx2.1-Cre (n=7) or FoxP2-Cre (n=9) animals, respectively.

For the behavioural locomotion experiments, opto-excitation experiments were performed in animals (D2-Cre, n = 7; Vglut2-Cre, n=7; FoxP2-Cre, n = 6) that were injected bilaterally into the region of interest (i.e. striatum, STN, or GP) using the same virus as the one used for electrophysiological experiments: an AAV5-EF1α-DIO-ChR2(H134R)-eYFP (for D2-Cre and Vglut2-Cre mice) or an AAV5-EF1α-DIO-ChR2(H134R)-mCherry (for FoxP2-cre) at similar coordinates and volume as aforementioned. To control for a light effect on locomotion, we additionally performed opto-control experiments in animals (D2-Cre, n = 2; Vglut2-Cre, n=2; FoxP2-Cre, n = 2) injected bilaterally (in similar target regions) with the control virus AAV5-EF1α-DIO-eYFP (6.5×10^12^ viral particles/mL). Then, two ceramic optic fibers (CFX128, ThorLab) were implanted bilaterally within the dorsolateral striatum (AP: 1.2 mm anterior to bregma, ML: 1.9 mm, DV: −2.0 mm) the STN (AP: −2.0 mm posterior to bregma, ML: 1.5 mm, DV: −3.9 mm,), and the GP (AP: 0.35 mm posterior to bregma, ML: 1.9 mm, DV: −2.9 mm).

### *In vivo* electrophysiological recording

Electrophysiological recordings and optogenetic manipulations were realized at least 3 weeks after viral transduction under urethane anaesthesia. Anaesthesia was first induced with isoflurane (3-5%; Iso-vet^®^, Piramal healthcare), and maintained with urethane (i.p., 1.3g/Kg.; CAT#U2500, Sigma-Aldrich). The animals were then secured in a stereotaxic frame (Unimécanique, M2e) and placed on heating blanket. After subcutaneous injection of xylocain, skull was exposed and craniotomies were performed to enable electrophysiological recordings and optogenetic stimulation of the region of interest (same stereotaxic coordinates as above). In each mice, an electrocorticogram (ECoG) of the sensorimotor cortex was realized (AP: 2.2 mm rostral to bregma, ML: 2.1 mm) with a 1 mm screw juxtaposed to the dura mater. All along the recording session, saline solution was used to prevent dehydration and anaesthesia level was frequently controlled by examination of EcoG and by testing the response of gentled sensory stimuli (tail pinch).

In vivo extracellular recordings were performed using glass electrodes (1-3μm tip end, 12-20MΩ, GC150F, WPI) filled with a chloride solution 0.5 M containing neurobiotin tracer (1-2%, w/v; CAT#SP-1120, Vector laboratories) to perform juxtacellular labelling. Optogenetic manipulation of the recorded neurons were performed using optical fibers placed in the striatum and/or the STN and an opto-electrode, homemade by gluing an optical fiber (multimode Fiber, 0.22NA, core diameter: 105μm, Thorlabs) 1mm from the tip of the glass electrodes. This method allows to combin extracellular recording, juxtacellular labelling, and optogenetic manipulation. Once opto-electrode was implanted in the recording structure, the signal was amplified by 10 fold in axoClamp 2B (in bridge mode; Molecular Devices). Then, recording signal was amplified 1000 fold and filtered with differential AC amplifier (spike unit: 0.3-10 kHz, local field potential: 0.1Hz-10 kHz; model 1700, A-M Systems). For ECoG recording, the signal was amplified 1000 fold and filtered with the same differential AC amplifier (0.1Hz-5 kHz). Finally, the recorded signals were digitalized at 20 kHz, by using the Power1401-3 connected to a computer equipped with Spike2 software (version 8, Cambridge Electronic Design). To identify the location of the recorded unit we performed juxtacellular labelling as previously described (Mallet et al., 2012). Then histological validation of the labelled neurons and the recording track of the electrode confirmed the recording locations. This approach has been used for all our experiments.

### *Ex vivo* electrophysiology recording

Mice with an age range of 56-105 postnatal day were sedated with isoflurane, deeply anesthetized with an i.p. injection of ketamine/xylazine (75/10 mg/Kg) cocktail and then perfused transcardially with cold (0-4°C) modified artificial cerebrospinal fluid (ACSF), continuously oxygenated with carbogen (95% O_2_ - 5% CO_2_) and containing the following (in mM): 230 sucrose, 26 NaHCO_3_, 2.5 KCl, 1.25 NaH_2_PO_4_, 0.5 CaCl_2_, 10 MgSO_4_ and 10 glucose (pH~7,35). The Brain was rapidly removed, glued to the stage of a vibratome (VT1200S; Leica Microsystems, Germany), immersed in the ice-cold ACSF and sectioned into 300 μm thick parasagittal or coronal slices. Slices containing the STR, GP and STN were transferred for 1 hour to a standard ACSF solution, warmed (35°C) and equilibrated with carbogen and containing (in mM unless otherwise stated): 126 NaCl, 26 NaHCO_3_, 2.5 KCl, 1.25 NaH_2_PO_4_, 2 CaCl_2_, 2 MgSO_4_, 10 glucose, 1 sodium pyruvate and 4.9 μM L-gluthathione reduced. Single slices were then transferred to a recording chamber, perfused continuously with equilibrated ACSF (without sodium pyruvate and L-gluthathione) heated at 32- 34°C, and visualized using infrared gradient contrast video microscopy (Ni-e workstation, Nikon) and a 60X water-immersion objective (Fluor APO 60X/1.00 W, Nikon). Recordings from individual GP neurons were performed with patch electrodes (impedance, 3–8 M*Ω*) fabricated from borosilicate glass capillaries (GC150F10; Warner Instruments, Hamden, CT, USA) pulled with a horizontal puller (P-97; Sutter Instruments, Novato, CA, USA). For whole-cell voltage-clamp recordings, pipettes were filled with K-Gluconate-based internal solution containing (in mM): 135 K-gluconate, 3.8 NaCl, 1 MgCl_2_.6H_2_O, 10 HEPES, 0.1 EGTA, 0.4 Na_2_GTP, 2 Mg_1.5_ATP, 5 QX-314 and 5.4 biocytin (pH=7.2, ~292 mOsm). In some recordings performed on D2-cre mice a cesium-based solution was used containing (in mM): 135 CsCl, 3.6 NaCl, 1 MgCl_2_.6H_2_O, 10 HEPES, 0.1 Na_4_EGTA, 0.4 Na_3_GTP, 2 Mg_1.5_ATP, 5 QX-314 and 5.4 biocytin (pH=7.2, ~292 mOsm). Recordings were obtained using a Multiclamp 700B amplifier and Digidata 1440A digitizer controlled by Clampex 10.3/10.6 (Molecular Devices LLC). Signals were sampled at 20 kHz and low-pass filtered at 4 kHz. Whole-cell voltage clamp recordings with CsCl and K-gluconate filled electrodes were corrected for a junction potential of −4 mV and −13 mV respectively. Series resistance was monitored throughout the experiment by voltage steps of −5 mV. Data were discarded when the series resistance changed by >20%. After electrophysiological recordings, slices were fixed overnight in a solution of paraformaldehyde at 4% and maintained in PBS-azide at 0.2% at 4°C until immunohistochemical processing.

### Drugs

Unless otherwise stated, all drugs were purchased from BioTechne and prepared in distilled water as concentrated stock solutions and stored at −20°C. On the day of the experiment, drugs were diluted and applied through the bath perfusion system. When necessary, GABAergic and glutamatergic antagonists were used to block the synaptic transmission (data not shown). GABA_A_ receptors were blocked with 20 μM 4-[6-imino-3-(4-methoxyphenyl) pyridazin-1-yl] butanoic acid hydrobromide (SR95531/GABAzine). NMDA receptors were blocked with 50 μM D-(–)-2-amino-5-phosphonopentanoic acid (D-APV). AMPA/kainate receptors with 20 μM 6,7-dinitroquinoxaline-2,3-dione (DNQX disodium salt).

### Optogenetic stimulation

For *in vivo* experiments, optogenetic stimulations were performed through optical fibers (multimode Fiber, 0.22NA, core diameter: 105μm, Thorlabs) implanted into the right striatum and/or STN (coordinates as above). The power at the tip of the optics fibers was always measured with a power meter (PM100D, Thorlabs) right before insertion in brain tissue. Two optogenetic stimulation protocols were used during the electrophysiological recordings in this study. The first one consisted of a 2 s long continuous blue light stimulation (0.5 mW measured at the tip of optic fiber) applied every 10sec. The 2 s preceding the light pulse was used as the baseline (i.e. OFF epoch) and was compared to the effect of the 2 s long optogenetic stimulation (i.e. ON epoch). Only neurons recorded for at least 20 laser stimulations were kept for further analysed. The second protocol consisted of a short 10 ms blue light stimulation (0.5 mW) applied every second. For behavioural experiments, optogenetic stimulation consisted of 500 ms blue light stimulation (1 mW) applied randomly with an inter-laser interval of at least 3 s. The laser bench (Errol laser) was controlled by analog signals sent by a Power1401-3 (Cambridge Electronic Design) connected to a computer equipped with Spike2 software. For *ex vivo* optogenetic stimulation, we used a LED laser source (Prizmatix, Israel) connected to an optic fiber (core diameter: 500 μm) placed directly above the brain slice. For cell body stimulation, we used continuous 1 s long light stimulation at low intensity (4 mW). To evoke reliable synaptic transmission, single pulses or train of stimulation (20 pulses at 20Hz) of 1ms duration at full power (90 mW) were used in order to maximize axon terminal depolarization and efficient release of neurotransmitter (Jackman et al., 2014).

### Juxtacellular labelling

In this study, we performed juxtacellular labelling to identify the location of the recorded unit as previously described (Mallet et al., 2012; Pinault, 1996). Briefly, the recorded unit was stimulated by a current pulse of 250ms long (50 % duty cycle, 1-10 nA) sent through the recording electrode by axoClamp 2B (in bridge mode; Molecular Devices). Then, the location and the molecular profile of the labelled neurons were determined by histological verification.

### Histology

At this end of recording day, the mice were sacrificed with an overdose of pentobarbital sodium (150 mg/Kg, i.p.; Axience). An intracardiac perfusion (PBS 0.01mM following by formaldehyde 4%) was then performed for further histological validations. The brain was kept overnight in a solution of PBS 0.01 mM/formaldehyde 4% (v/v; CAT#20909.330, VWR) and then cut in 50 μm slices with a vibratome (VT1000, Leica Microsystemes). To reveal the juxtacellularly labelled neurons, the slices were incubated overnight in a solution of PBS 0.01 mM/Triton™ X100 0.3% (v/v; CAT#T9284, Sigma-Aldrich) containing streptavidine-CY3 Zymax (1/1000, v/v; CAT#438315, Life Technologies). The slices were then washed in PBS 0.01 mM before mounting onto slides in Fluoromount medium (Cliniscience). Additional staining was performed through indirect immunofluorescence staining using primary and fluorescent secondary antibodies. For all immunostainings, the slices were incubated overnight in a solution of PBS 0.01mM / Triton™ X100 0.3% (v/v) containing the primary antibody. The slices were then washed 3 times in PBS 0.01 mM, incubated 4h in a solution of PBS 0.01 mM / Triton™ X100 0.3% (v/v) containing the secondary antibody, washed again in PBS 0.01 mM and mounted onto slides in Fluoromount medium. The primary antibodies used in this study were: the rabbit anti-Nkx2.1 (or anti-TTF-1, 1/500, v/v; CAT#H-190, Santa Cruz), the rabbit anti-preproenkephalin (1:5000, LifeSpan Biosciences, LS-C23084), and goat anti-FoxP2 (1/500, v/v; CAT#N-16, Santa Cruz). The immunolabelling for the preproenkephalin was optimized using the heat-treatment antigen retrieval method as described previously (Mallet et al., 2012). The secondary antibodies used were: Alexa Fluor^®^ 488 donkey anti-chicken (1/500, v/v; CAT#703-545-155, Jackson ImmunoResearch), CY™5-conjugated donkey anti-rabbit (1/500, v/v; CAT#711-175-152, Jackson ImmunoResearch), and Brilliant Violet™421-conjugated donkey anti-goat (1/500, v/v; CAT#705-675-147, Jackson ImmunoResearch).

To reveal biocytin-filled neurons after *ex vivo* recordings, brain slices were incubated overnight in a solution of PBS 0.01 mM/Triton™ X100 0.3% (v/v; CAT#T9284, Sigma-Aldrich) containing streptavidine-488, (1/500; Cat#S11223; Life technologies), or streptavidine-557 (1/1000; Cat# NL999, R&D system). Additional staining was performed as for the *in vivo* histology (see above), but with only 2 hours of incubation with the secondary antibody. The primary antibodies used in this study were: the rabbit anti-Nkx2.1 (same ref as above), goat anti-FoxP2 (same ref as above), Rabbit anti-RFP (1/1000; Cat# R10367, Life technologies), Rat anti-RFP (1/500, v/v; Cat#5f8, Chromotek), Mouse anti-GFP (1/600; Cat#11814460001, Roche). The secondary antibodies used were: Alexa-Fluor Donkey anti-Rabbit 488 (1/500, v/v; Cat#A21206, Life technologies), Alexa-Fluor Donkey anti-Mouse 488 (1/500, v/v; CAT#A 21202, Life technologies), Alexa-Fluor Donkey anti-Rabbit 568 (1/500 v/v; Cat#A11075, Life technologies), Alexa-Fluor Donkey anti-Rat 568 (1/500 v/v; Cat#A11077, Invitrogen), Alexa-Fluor Donkey anti-Rat 594 (1/500 v/v; Cat# A21209, Life technologies), Alexa-Fluor Donkey anti-Goat 647 (1/500; Cat#A21447, Life technologies). The slices were then washed in PBS 0.01 mM before mounting onto slides in vectashield medium (CAT#H-1000, Vector laboratories). The low magnification images were acquired with an epifluorescence microscope (Axio Imager 2, Zeiss). The high magnification images were acquired with a confocal microscope (Leica TCS SP8) head mounted on an upright stand DM6 FS (Leica Microsystems, Mannheim, Germany) with a HC Plan Apo CS2 63X oil NA 1.40 objective and equipped with a motorized XY stage, and a galvanometric stage for fast Z acquisition. The confocal is equipped with four laser lines (405 nm, 488 nm, 552 nm and 638 nm), a conventional scanner (10-1800 Hz), and detectors (two internal PMT and two internal hybrid detectors). The images were further extracted using Fidji (Schindelin et al., 2012).

### Data analysis of electrophysiological recordings

*In vivo recordings:* All data processing (spike sorting and ECoG filtering) were realized offline with Spike2 software (Cambridge Electronic Design). Peri-stimulus time histograms (PSTH) of each spike trains were generated using spike2 and a custom Matlab^®^ script for the 2 s long (width: 6 s, offset: 2 s, bin size: 50 ms) and the 10 ms light stimulation protocol (width: 0.180 s, offset: 0.03 s, bin size: 2.5 ms). A response was classified as statistically significant if the spike count values of 3 consecutive bins of light pulse delivery were < −2 standard deviation (SD) for inhibitory responses, or > 2 SD for excitatory responses. SD was measured during the baseline OFF epoch. We also analysed in spike2 the frequency and the coefficient of variation (CV) of each spike train during OFF and ON epoch. *Ex vivo recordings:* Ex vivo electrophysiological data were analyzed using ClampFit 10.3/10.6 (Molecular Devices LLC) and OriginPro (version 7 and 9; OriginLab Corporation). Spontaneous firing rate and standard deviation of interval interspike (SDisi) of GP neurons were calculated by detection of action potentials over a period of 100-150 s (Deister et al., 2013) in ClampFit 10.3/10.6. PSTH of each spike trains for continuous 1 s or pulse train (20 pulses, 20Hz) optogenetic stimulations were generated using OriginPro with a bin size of 50 ms. Both frequency and count PSTH were generated for each spike train to evaluate the effect of the optogenetic manipulations. The magnitude of light-evoked inhibitory post-synaptic currents (IPSC) or excitatory post-synaptic currents (EPSC) were obtained by averaging 6-10 trials. Currents were then converted into conductance according to the formula (g=i/(V_rec_-E_NT_) were V_rec_ corresponds to the membrane potential of the recording and E_NT_ the reversal potential for GABA or Glutamate.

### Behavioural locomotion experiments

Mice were daily trained to walk continuously for 5 minutes on a motorized treadmill belt (BiosebLab) running at 15cm/s for at least one week before the beginning of optogenetic experiments. During the experiments, the locomotor sequences were video capture (250 frames/s) bilaterally using a pair of high speed cameras (Basler ac 1920-150uc, Basler AG) mounted with Fujinon lenses (model HF16XA-5M) and synchronized using a trigger cable with the software Streampix (Norpix, Inc). Video recordings were stored on a computer and analysed off-line.

### Data analysis of locomotion

The video frames corresponding to period of laser stimulation were extracted using home-made routines based on the OpenCV-python library. The locomotor activity of the animal measured on the treadmill was obtained from a frame by frame analysis of body landmarks position (base of the tail, fore- and hindlimb toes, and nose extremity). The X-Y coordinates of the landmarks were extracted using Deep Lab Cut (Mathis et al., 2018) on a linux workstation equipped with a NVIDIA GeForce RTX 2080TI graphic card. Before any further data processing, the Deep Lab Cut labelling output was then checked for errors of detections and corrected by first comparing the detections from the left and right camera frames. In addition, a second step of correction included the replacement of labelling that yielded to an instantaneous velocity above 28 cm/s (measured as being the maximal velocity reachable by the mouse in our conditions) by the averaged value measured in the time window covering 120 ms before and after the misdetection. Two of the foretold body landmarks (Nose extremity and Tail base) were chosen for quantification purposes. The velocity corresponding to each frame (accounting for 4 ms of the movement) was calculated using the displacement along the treadmill axis (averaged over the right and left labels and body landmarks) starting 60 ms before and finishing 60 ms after. This particular 120 ms window size in addition to a moving average of window size 20 ms that was performed after the velocity calculation assured minimizing the small velocity fluctuations captured in the measurements. The mean velocity was calculated for all the frames in a 500 ms time window preceding and following the laser onset. The average velocity averaged over the laser-OFF and laser-ON period has been reported as representations of the velocity in these periods.

### Statistical analysis

Experimental data were analysed using the computer programme GraphPad Prism (v8.4.3, GraphPad Software, Inc). Data in the figures are presented as the median and interquartile range unless otherwise stated. Box plots (central line: median; box: 25%-75%; whiskers: min-max) were used to illustrate sample distributions with individual values represented by circles or individual lines. Group data in the text and the supplementary tables S1-S14 are presented as mean ± standard error of the mean (SEM). For dependent sample, the paired *t*-test was used excepts if the normality distribution test failed (Shapiro-Wilk test, *p*<0.05). In the latter case, the Wilcoxon Signed Rank (WSR) test was used. For independent samples, we applied the normality (Shapiro-Wilk test) and equal variance tests. A Mann-Whitney U (MWU) test was used when the distributions were normal and the group variances were equal. Contingency was assessed using Fisher‘s exact test. For multiple repeated measured group comparisons, the Friedman test was used, followed by Dunn‘s post hoc test. For behavioural analysis, two-tailed, non-parametric statistical comparisons were made using Wilcoxon Signed Rank test (WSR) with Bonferroni correction to compare the OFF *vs.* ON average velocities measured within each animal group. Furthermore, the non-parametric Mann–Whitney U (MWU) test for unpaired data was used to compare the average laser-ON velocities between the ChR2 and the eYFP control groups. The level of statistical significance was set at *p* < 0.05.

## Supporting information

Supplemental data informations

## Acknowledgements

This work was supported by grants from the French Agence Nationale de la Recherche (ANR-14-CE13-0024-01 and ANR-15-CE37-0006), from the LABEX BRAIN (ANR-10-LABX-43), and the Association France Parkinson (grant # OPE-2018-0459). The microscopy was done in the Bordeaux Imaging Center, a service unit of the CNRS-INSERM and Bordeaux University, member of the national infrastructure France BioImaging supported by the French National Research Agency (ANR-10-INBS-04). We also thank the PIV-EXPE of the University of Bordeaux and the Ani*Motion* collaborative facility (INCIA). University of Bordeaux and CNRS provided infrastructural support. We are grateful to E. Doudnikoff, Monika Fernández-Monreal, A. Fayoux, T. Nguyen and H. Orignac for technical assistance as well as Drs. R. Schmidt, A. Leblois, G. Silberberg,, F. Georges, M. Deffains, and R. Bidgood for insightful scientific discussions and comments on the manuscript. The authors declare no competing financial interests.

## Author Contributions

J. B. and N. P. M. conceptualized the project. A. A., B. C., L.G., S. G. and N. P. M. performed *in vivo* electrophysiology recordings. M. B. and J. B. performed slice electrophysiology recordings. A. A., S. F. M. B. and N. P. M. performed histology. A. A., S. A. L., G. B. and N. P. M. performed behavioural experiments. A. A., M. B., J. B., S. A. L., G. C. and N. P. M. performed the analysis. J.B. and N. P. M. designed the experiments and supervised the project. A. A., M. B., J.B and N. P. M. wrote the manuscript.

