## Supplemental data informations for "A disynaptic circuit in the globus pallidus controls locomotion inhibition"

Figure S1

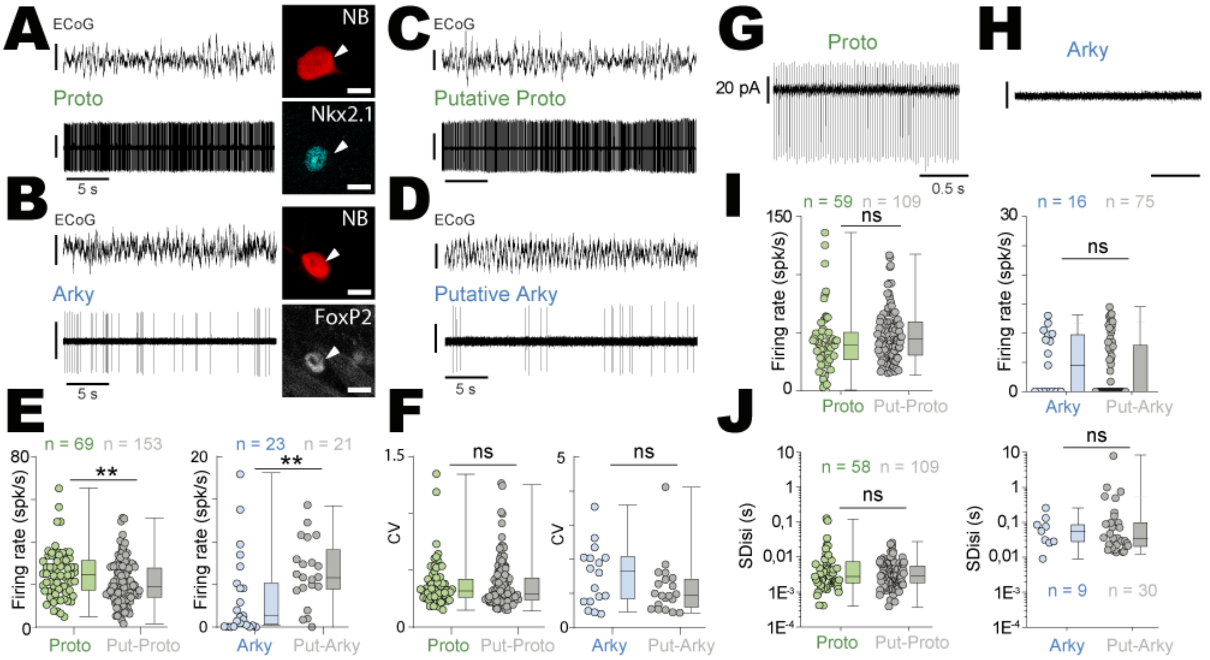


**Figure S1 *In vivo* and *ex vivo* electrophysiological properties of labelled *vs.* putative prototypic and arkypallidal neurons.** ***(A, B)*** Raw *in vivo* electrophysiological traces of identified prototypic ***(A)*** and arkypallidal ***(B)*** GP neurons recorded simultaneously with the ECoG activity. Neuronal identity was confirmed with juxtacellular neurobiotin labelling and immunodetection for the expression of Nkx2.1 for prototypic (top) or FoxP2 for arkypallidal neurons (bottom), scale bars: 10$\mu$m. ***(C, D)*** Raw *in vivo* electrophysiological recordings of putative prototypic ***(C)*** and arkypallidal GP ***(D)*** neurons with the corresponding ECoG activity. ***(E, F)*** Comparison of the *in vivo* firing rate ***(E)*** and coefficient of variation (CV, ***F)*** for juxtacellularly labelled vs. unidentified prototypic (left) and arkypallidal (right) GP neurons recorded *in vivo*. ***(G, H)*** Raw *ex vivo* spontaneous activity of identified prototypic ***(G)*** and arkypallidal ***(H)*** GP neurons. ***(I, J)*** Comparison of the *ex vivo* firing rate ***(I)*** and interspike interval standard deviation ($SD_{ISI}$, ***J***) for identified vs. putative prototypic (left) and arkypallidal (right) neurons. ** p<0.01, ns: not significant. See Table S7 for more details and statistical information.

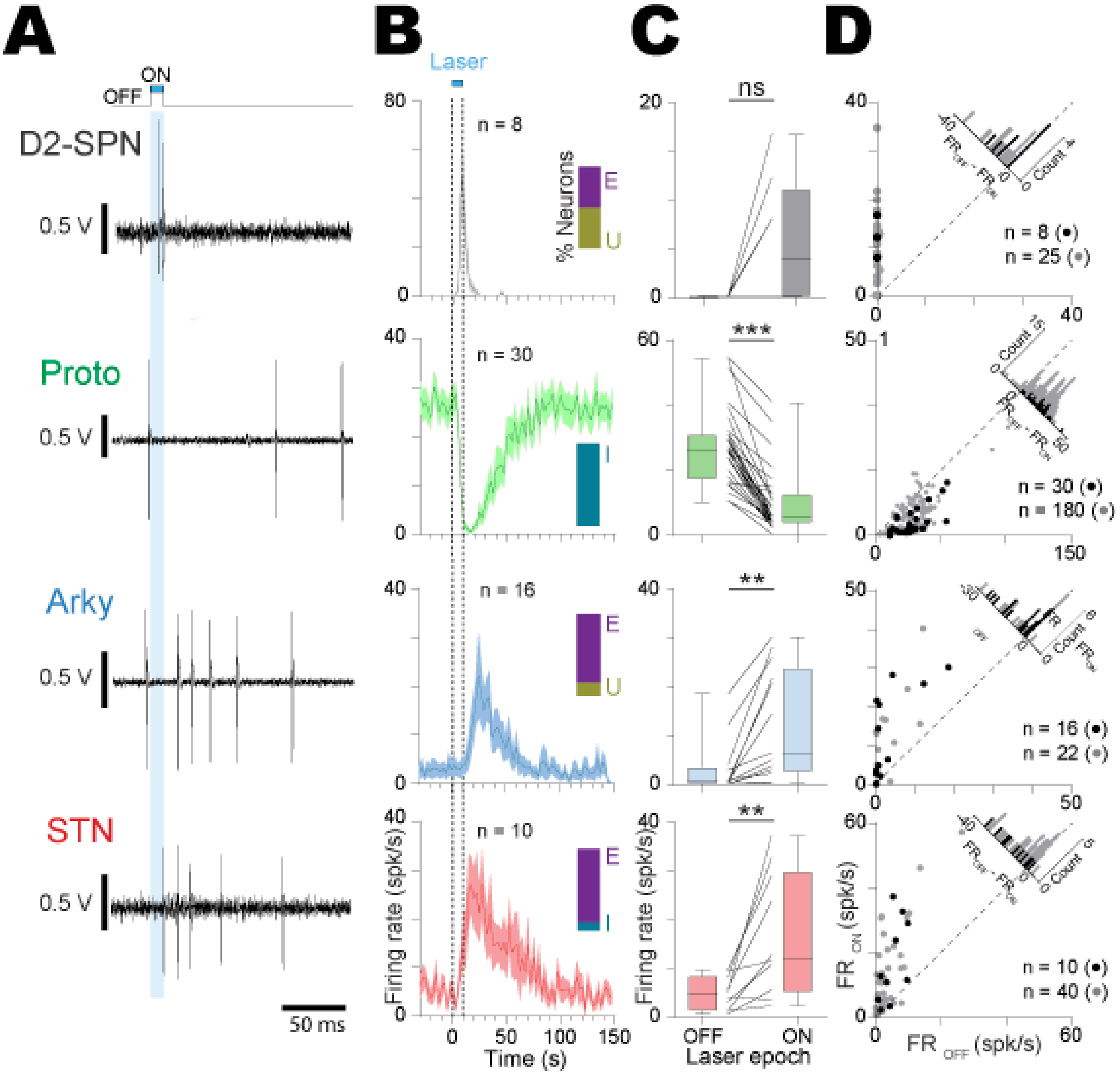


**Figure S2 Short-duration opto-activation of D2-SPNs induced opposing effects on prototypic and arkypallidal neurons.** ***(A)*** Typical examples of *in vivo* single-unit activity for one juxtacellularly labelled D2-SPN, prototypic, arkypallidal, and STN neurons during D2-SPN opto-activation (10 ms blue light pulses). ***(B)*** Population PSTH (bin size: 2.5 ms) of all the juxtacellularly labelled D2-SPN (n= 8, grey), prototypic (n= 30, green), arkypallidal (n= 16, blue), and STN neurons (n= 10, red) during opto-activation of D2-SPN. ***(C)*** Box-and-whisker plots representing the average firing rate of all the labelled D2-SPNs (grey), prototypic (green), arkypallidal (blue), and STN neurons (red) during OFF *vs.* ON laser stimulation of D2-SPNs. ***(D)*** Scatter plots representation of juxtacellularly labelled (black dots) and putative (grey dots) D2-SPN, prototypic, arkypallidal, and STN neurons during D2-SPN ON/OFF opto-stimulation. Group data represents mean ± SEM, box-and-whisker plots indicate median, first and third quartile, min and max values. ** p<0.01, *** p<0.001, ns: not significant. See Table S8 for more details and statistical information.

Figure S3

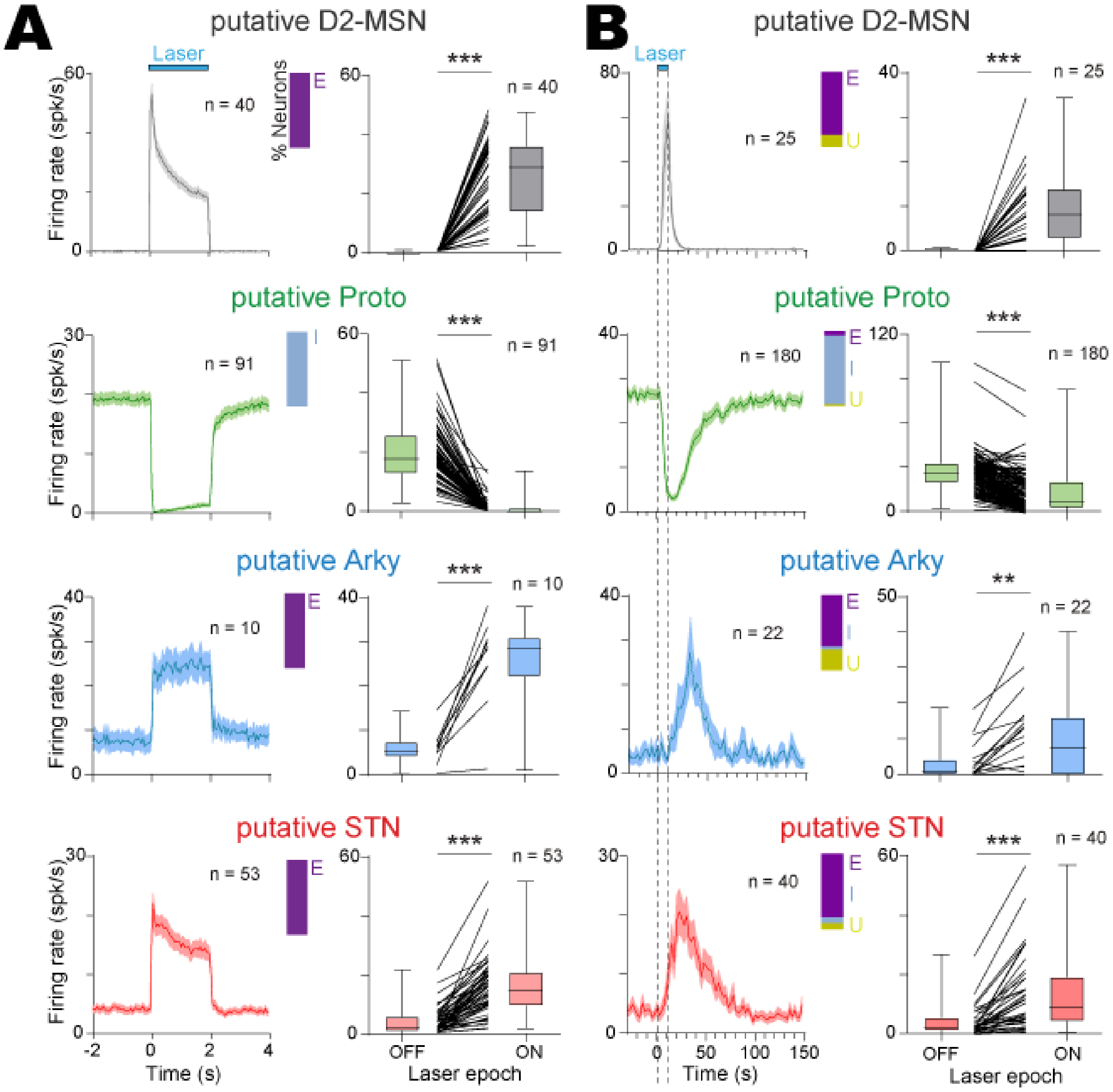


**Figure S3 Opto-activation of D2-SPNs induced opposite effects in putative prototypic and putative arkypallidal neurons.** ***(A, B)*** Population PSTH (left) and average firing rate responses (right) calculated during a 2s long (***A***, bin size: 50 ms) or a 10 ms short (***B***, bin size: 2.5 ms) opto-excitation of D2-SPNs for putative D2-SPN (grey), prototypic (green), arkypallidal (blue), and STN neurons (red). Group data represents mean ± SEM, box-and-whisker plots indicate median, first and third quartile, min and max values. ** p<0.01, *** p<0.001. See Table S9 for more details and statistical information.

Figure S4

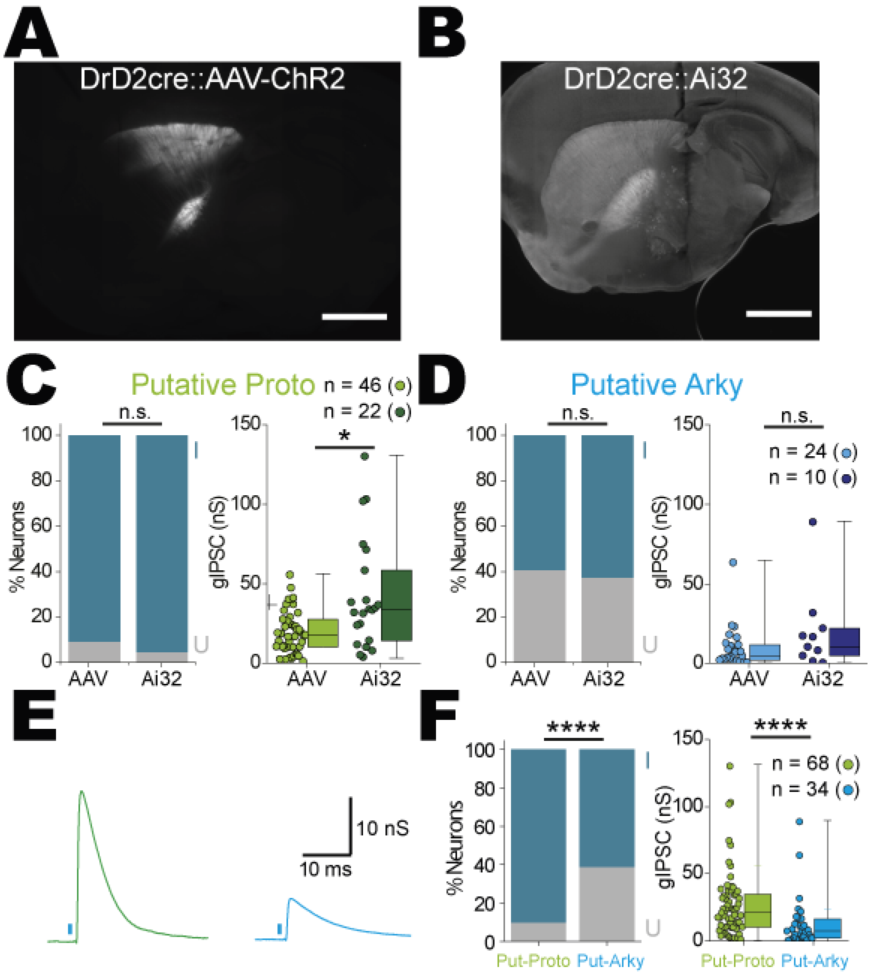


**Figure S4** ***Ex vivo*** **characterization of D2-SPN inputs onto putative GP neurons.** ***(A, B)*** Epifluorescent images illustrating the virus-induced ChR2-eYFP expression in sub-population of D2-SPNs (***A***) or the constitutive expression of ChR2-eYFP using D2-Cre::Ai32 mice that labelled all D2-SPNs (***B***) in the striatum and their respective projections to the GP (scale: 0.5 mm). ***(C)*** Bar graph representing the percentage of IPSC-recipient (right) and the averaged IPSC conductance (left) for all the putative prototypic neurons recorded in D2-Cre::AAV-ChR2-eYFP mice (AAV and light green) or in D2-Cre::Ai32 mice (Ai32 and dark green). ***(D)*** Bar graph representing the percentage of IPSC-recipient (right) and the averaged IPSC conductance (left) for all the putative arkypallidal neurons recorded in D2-Cre::AAV-ChR2-eYFP mice (AAV and light blue) or in D2-Cre::Ai32 mice (Ai32 and dark blue). ***(E)*** Typical inhibitory post-synaptic currents (IPSCs) recorded in putative prototypic (green) or arkypallidal (blue) neurons during the laser-stimulation (1ms pulse) of D2-SPN axon terminals in D2-Cre mice injected with an AAV-ChR2 in the striatum. ***(F)*** Bar graph representing the percentage of IPSC-recipient (left) and the averaged magnitude of IPSC conductance (right) measured in putative prototypic and arkypallidal neurons. * $p<0.05$, **** $p<0.0001$, ns: not significant. See Table S10 for more details and statistical information.

Figure S5

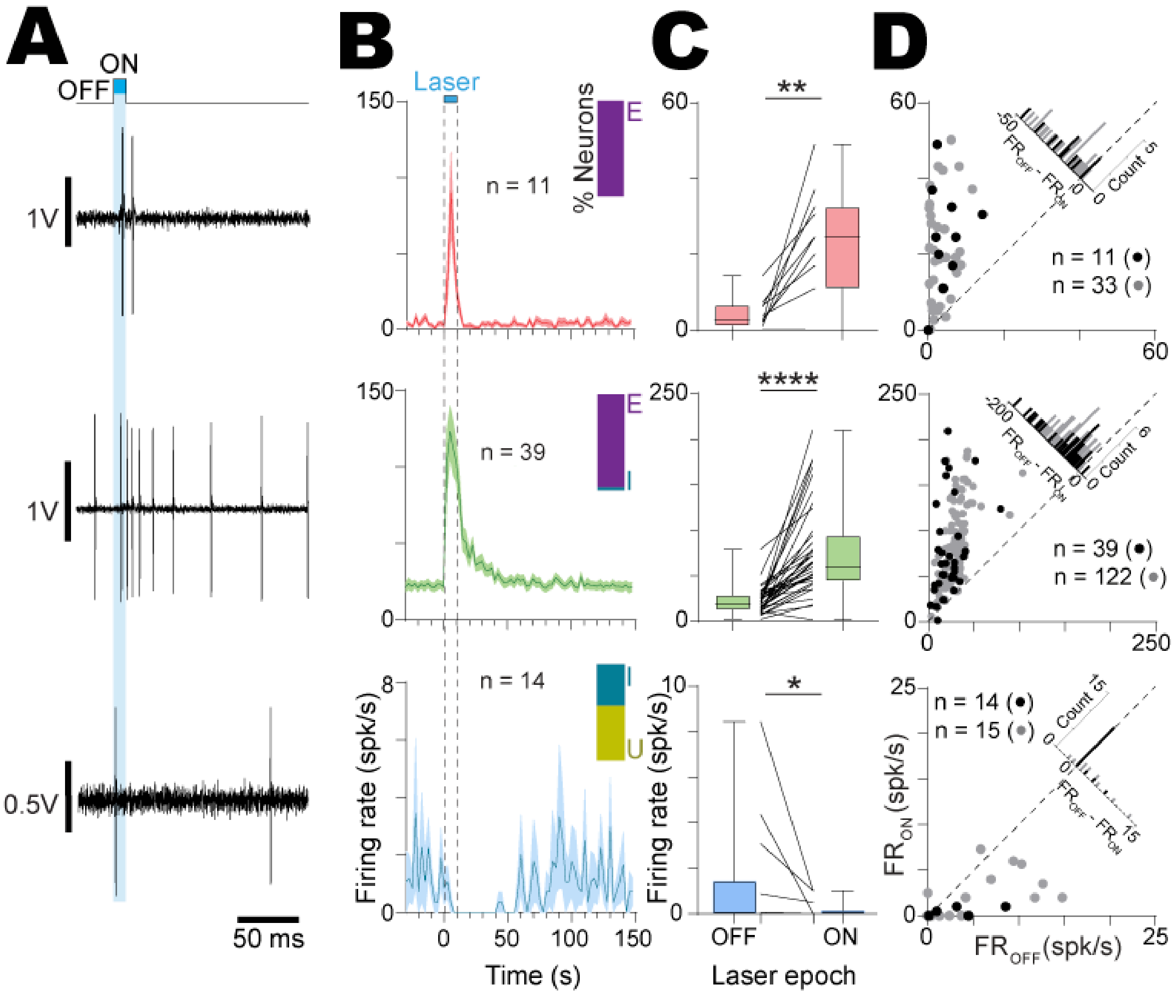


**Figure S5 Short-duration opto-activation of STN neurons induced opposite effects on prototypic and arkypallidal neurons *in vivo*.** ***(A)*** Representative examples of *in vivo* single-unit activity for one juxtacellularly labelled STN, prototypic, and arkypallidal neurons during STN opto-activation (10 ms blue light pulses). ***(B)*** Population PSTH (bin size: 2.5 ms) of all juxtacellularly labelled STN (n= 11), prototypic (n= 39) and arkypallidal neurons (n= 14) during opto-activation of STN neurons. The inset bar plot represents the percentage of neurons excited (E), inhibited (I) and unaffected (U) by the light stimulation. ***(C)*** Box-and-whisker plots representing the average firing rate of all the labelled STN (red), prototypic (green) and arkypallidal (blue) neurons during OFF *vs.* ON laser stimulation of STN neurons. ***(D)*** Scatter plots representation of juxtacellularly labelled (black dots) and putative (grey dots) STN, prototypic and arkypallidal neurons during STN neurons ON/OFF opto-stimulation. Group data represents mean ± SEM, box-and-whisker plots indicate median, first and third quartile, min and max values.* $p<0.05$, ** $p<0.01$, **** $p<0.0001$. See Table S11 for more details and statistical information.

Figure S6

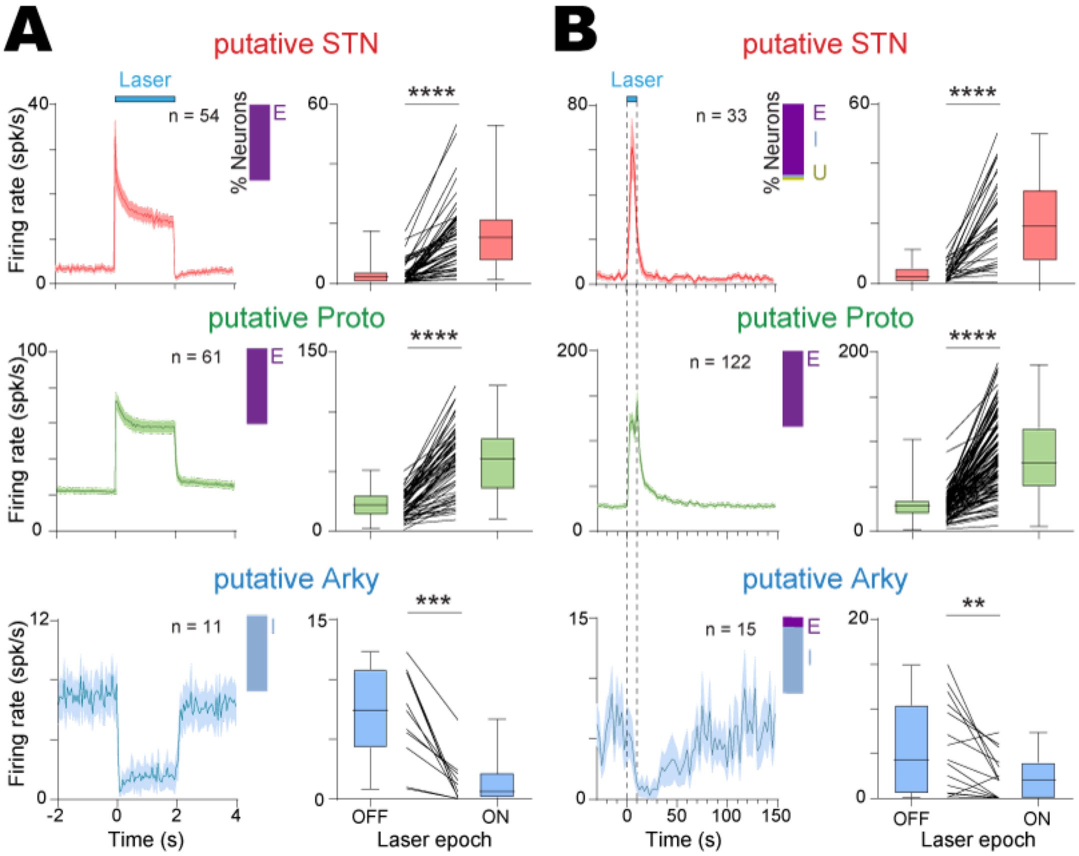


**Figure S6 Opto-activation of STN neurons induced opposite effects on putative prototypic and putative arkypallidal neurons *in vivo*. *(A, B)*** Population PSTH (left) and averaged firing rate responses (right) calculated during a 2s long (***A***, bin size: 50 ms) or a 10 ms short (***B***, bin size: 2.5 ms) opto-excitation of STN neurons for putative STN (red), prototypic (green), arkypallidal (blue) neurons. The inset bar plots represent the percentage of neurons excited (E), inhibited (I) an unaffected (U) by the light stimulation. Group data represents mean ± SEM, box-and-whisker plots indicate median, first and third quartile, min and max values. ** p<0.01, *** p<0.001, **** p<0.001. See Table S12 for more details and statistical information.

Figure S7

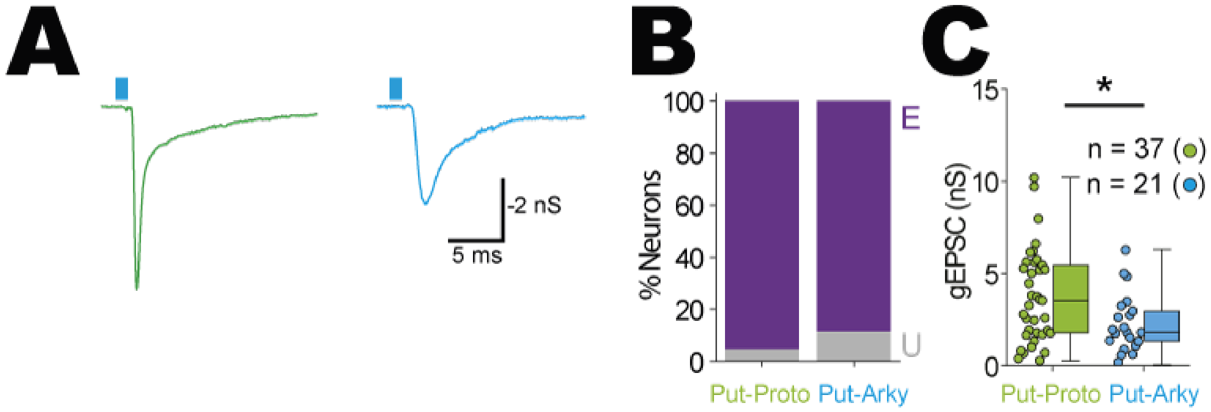


**Figure S7 Opto-activation of STN inputs induced excitatory post-synaptic currents in putative GP neurons *ex vivo*. *(A)*** Representative excitatory post-synaptic currents (EPSCs) evoked by light-activation (1ms) of STN axon terminals of Vglut2-Cre mice injected in the STN with an AAV-DIO-ChR2. ***(B)*** Bar graph representing the percentage of EPSC-recipient (purple) putative prototypic and arkypallidal neurons. E: excited; U: unchanged. ***(C)*** Population data depicting the greater magnitude of EPSC conductance in putative prototypic compared to arkypallidal neurons. * $p<0.05$. See Table S13 for more details and statistical information.

Figure S8

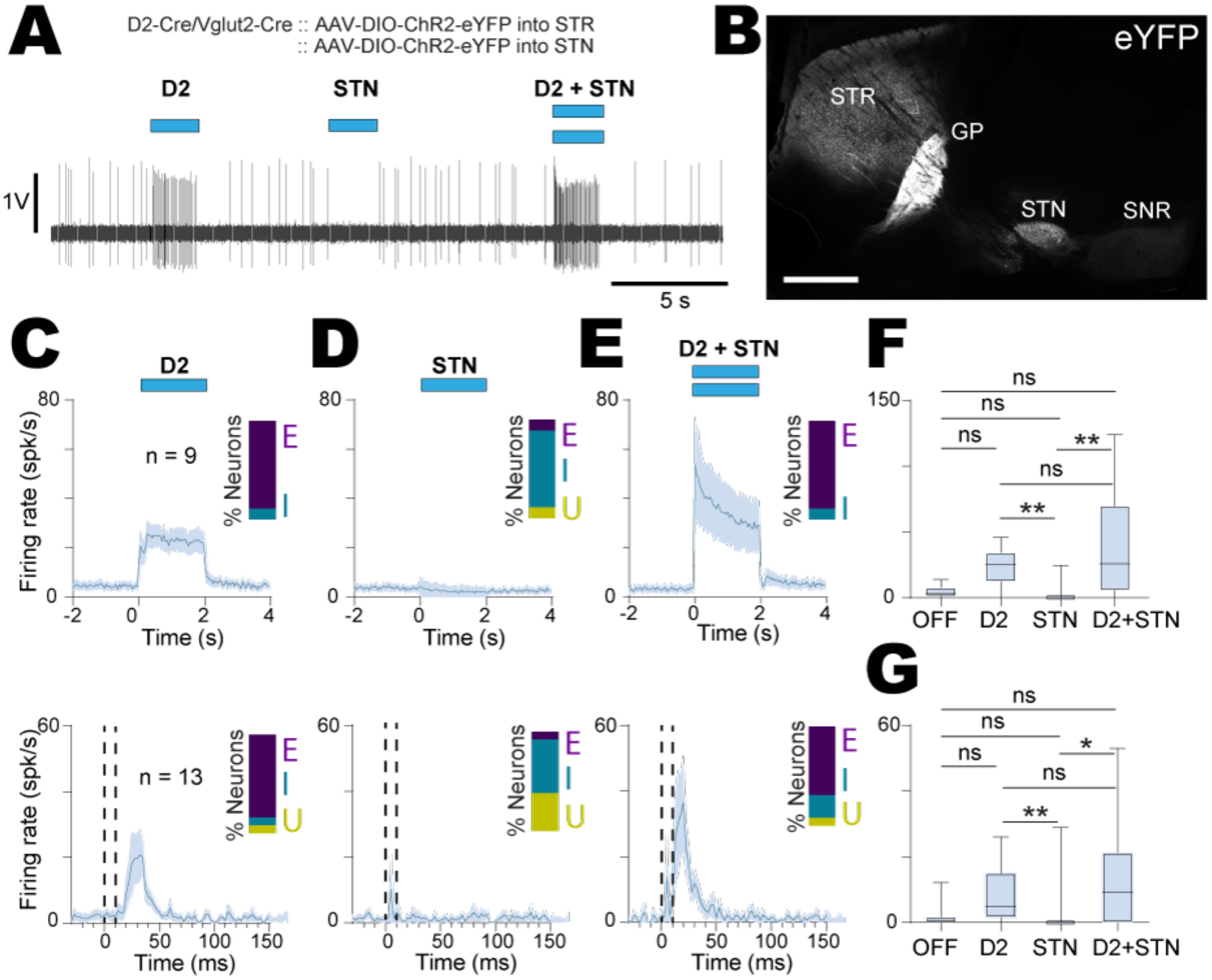


**Figure S8 The *in vivo* integration of STN inputs in arkypallidal neurons are gated by the presence of D2-SPNs inputs.** ***(A)*** Representative single-unit recording showing the effect of D2-SPN (D2 stim), STN (STN stim) and D2-SPN + STN (D2 stim + STN stim) opto-activation (2 s blue light pulses) on the firing activity of one arkypallidal neuron. ***(B)*** Epifluorescent image illustrating the ChR2-eYFP expression in D2-SPN and STN neurons with dense axonal projections to the GP (scale: 1 mm). ***(C, D, E)*** Population PSTH of arkypallidal neurons calculated during a 2s long (top, n = 9, bin size: 50 ms) or a 10 ms short (bottom, n = 13, bin size: 2.5 ms) opto-activation of D2-SPN ***(C)***, STN ***(D)*** and D2-SPN + STN ***(E)*** inputs. The inset bar plots represent the percentage of neurons excited (E), inhibited (I) and unaffected (U) by the light stimulation. ***(F, G)*** Average firing rate changes in arkypallidal neurons calculated during a 2s long (***F***) or a 10 ms short (***G***) OFF *vs*. ON opto-activation of D2-SPN (D2), STN (STN) and D2-SPN + STN (D2+STN) inputs. Group data represents mean ± SEM, box-and-whisker plots indicate median, first and third quartile, min and max values. * p<0.05, ** p<0.01, ns: not significant. See Table S14 for more details and statistical information.

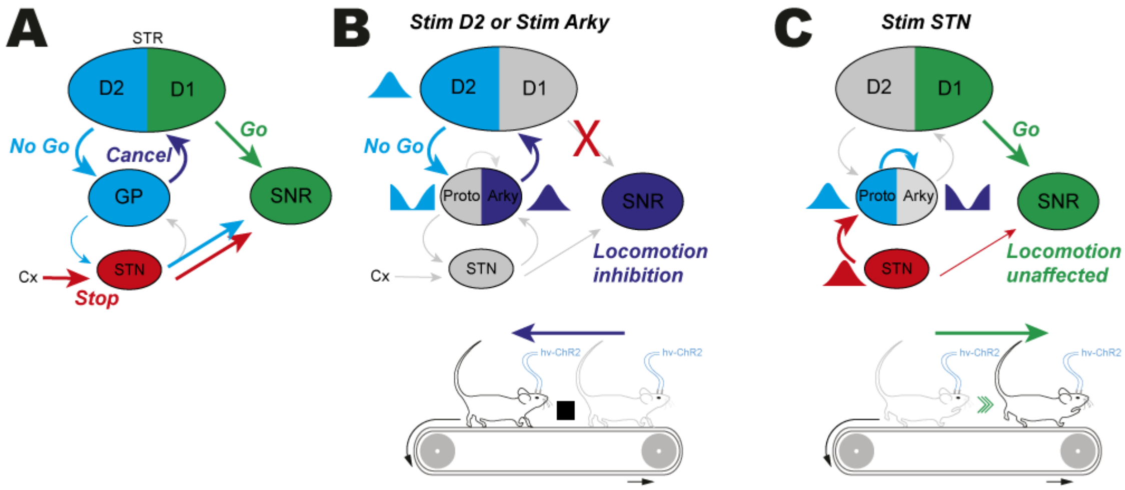


**Figure S9 An intra-GP circuit for the control of locomotion inhibition.** *(A)* Hypothetical basal ganglia multiple pathways to control locomotion. ***(B)*** Activation of the indirect pathway (D2) suppresses activity in prototypic (Proto) neurons which turns ON arkypallidal (Arky) neurons and inhibits locomotion. ***(C)*** Stimulation of the hyperdirect pathway (STN) produces the opposite effect on prototypic and arkypallidal neuron activity but fails to inhibit movement. Cx: cortex; STR: striatum.

**Supplementary tables:**

***Table S1. Relates to Figure 1***

| Figure | Parameter | n (mice/neurons) | Data Type | Data Value (mean ± SEM) | Statistical test | significance level |
|---|---|---|---|---|---|---|
| **1D** | D2-SPNs Firing rate | (4/8) | OFF | 0,081 ± 0,058 | Wilcoxon signed rank test | *p*=0,0078 |
| | | | ON | 31,88 ± 3,11 | | |
| **1E** | Proto Firing rate | (24/36) | OFF | 26,12 ± 1,70 | Paired t test | t=13,79, *p*<0,0001 |
| | | | ON | 1,02 ± 0,34 | | |
| **1F** | Arky Firing rate | (13/14) | OFF | 4,41 ± 1,51 | Wilcoxon signed rank test | *p*=0,0002 |
| | | | ON | 20,83 ± 3,28 | | |
| **1G** | STN Firing rate | (5/10) | OFF | 4,92 ± 0,98 | Paired t test | t=7,036, *p*<0,0001 |
| | | | ON | 19,21 ± 2,39 | | |
| **1J** | Firing rate | (30/59) | Proto | 42.51 ± 3.48 | Mann-Whitney U test | *U=44.5, p*<0.0001 |
| | | (14/16) | Arky | 5.2 ± 1.27 | | |
| | SDisi | (29/58) | Proto | 0.0109 ± 0.0034 | Mann-Whitney U test | *U=32, p*<0.0001 |
| | | (9/9) | Arky | 0.0775 ± 0.0256 | | |
| **1L** | Input-recipient D2-SPN | (22/57) | % of Proto | 94.74 % | *Fischer exact test* | *p*=0.0146 |
| | | (12/12) | % of Arky | 66.67 % | | |
| **1L** | Conductance | (28/54) | Proto | 24.28 ± 2.49 | Mann-Whitney U test | *U=18, p*<0.0001 |
| | | (8/8) | Arky | 3.487 ± 1.08 | | |

*Table S2. Relates to Figure 2*

| Figure | Parameter | n (mice/neurons) | Data Type | Data Value (mean ± SEM) | Statistical test | significance level |
|---|---|---|---|---|---|---|
| **2D** | STN Firing rate | (8/74) | OFF | 6,12 ± 0,55 | Wilcoxon signed rank test | *p*<0,0001 |
| | | | ON | 1,11 ± 0,23 | | |
| **2I** | Proto Firing rate | (10/25) | OFF | 28,18 ± 1,79 | Friedman test with Dunn's multiple comparisons | Off *vs.* D2: Z=6.408, *p*<0.0001<br>Off *vs.* STN: Z=3.232, *p*=0.0074<br>Off *vs.* D2+STN: Z=6.792, *p*<0.0001<br>D2 *vs.* STN: Z=3.177, *p*=0.0089<br>D2 *vs.* D2+STN: Z=0.3834, *p*>0.9999<br>STN *vs.* D2+STN: Z=3.560, *p*=0.0022 |
| | | | D2 | 0,57 ± 0,26 | | |
| | | | STN | 10,29 ± 1,52 | | |
| | | | D2 + STN | 0,44 ± 0,21 | | |
| **2J** | Arky Firing rate | (7/9) | OFF | 3,31 ± 1,62 | Friedman test with Dunn's multiple comparisons | Off *vs.* D2: Z=3.651, *p*=0.0016<br>Off *vs.* STN: Z=1.643, *p*=0.6021<br>Off *vs.* D2+STN: Z=3.834, *p*=0.0008<br>D2 *vs.* STN: Z=2.008, *p*=0.2677<br>D2 *vs.* D2+STN: Z=0.1826, *p*>0.9999<br>STN *vs.* D2+STN: Z=2.191, *p*=0.1708 |
| | | | D2 | 24,50 ± 4,51 | | |
| | | | STN | 9,29 ± 2,26 | | |
| | | | D2 + STN | 20,95 ± 2,60 | | |

*Table S3. Relates to Figure 3*

| Figure | Parameter | n (mice/neurons) | Data Type | **Data Value** (mean ± SEM) | Statistical test | significance level |
|---|---|---|---|---|---|---|
| **3E** | STN Firing rate | (5/11) | OFF | 3,63 ± 1,26 | Wilcoxon signed rank test | *p*=0,001 |
| | | | ON | 16,12 ± 2,44 | | |
| | Proto Firing rate | (21/32) | OFF | 23,68 ± 2,23 | Wilcoxon signed rank test | *p*<0,0001 |
| | | | ON | 75,56 ± 6,98 | | |
| | Arky Firing rate | (9/9) | OFF | 2,16 ± 1,07 | Wilcoxon signed rank test | *p*=0,0078 |
| | | | ON | 0,16 ± 0,15 | | |
| **3J** | Input-recipient STN | (4/16) | % of Proto-GP | 100% | *Fischer exact test* | *p*=1 |
| | | (7/9) | % of Arky-GP | 100% | | |
| **3J** | Conductance | (4/16) | Proto-GP | 5.429 ± 0.81 | Mann-Whitney U test | *U=11,* *p*=0.0002 |
| | | (7/9) | Arky-GP | 1.423 ± 0.45 | | |

***Table S4. Relates to Figure 4***

| Figure | Parameter | n (mice/neurons) | Data Type | Data Value (mean ± SEM) | Statistical test | significance level |
|---|---|---|---|---|---|---|
| 4E | Proto Firing rate | (5/12) | OFF | 17.81 ± 5.61 | Wilcoxson Signed Rank | *p*=0.0005 |
| | | | ON | 97.14 ± 17.10 | | |
| 4F | Arky Firing rate | (5/8) | OFF | 10.28 ± 1.46 | Wilcoxson Signed Rank | *p*=0.0078 |
| | | | ON | 3.34 ± 0.97 | | |
| 4G | Conductance | (4/9) | Proto-GP | 6.17 ± 1.5 | Mann-Whitney U test | *p*=0.2775 |
| | | (4/10) | Arky-GP | 4.38 ± 1.07 | | |

***Table S5. Relates to Figure 5***

| **Figure** | **Parameter** | **n (mice/neurons)** | **Data Type** | **Data Value** (mean ± SEM) | **Statistical test** | **significance level** |
|---|---|---|---|---|---|---|
| 5E | Arky Firing rate | (6/10) | OFF | 11.86 ± 4.15 | Wilcoxson Signed Rank | *p*=0.0002 |
| | | | ON | 82.71 ± 14.93 | | |
| 5F | Proto Firing rate | (7/25) | OFF | 40.18 ± 4.41 | Wilcoxson Signed Rank | *p*=0.523 |
| | | | ON | 41.14 ± 4.38 | | |
| 5I | Proto Firing rate | (3/61) | OFF | 28.256 ± 1.290 | Paired *t* test | t=0.1571, *p*=0.8757 |
| | | | ON | 28.049 ± 1.981 | | |

***Table S6. Relates to Figure 6***

| Figure | Parameter | n (mice/laser stim) | Data Type | Data Value | Statistical test | significance level |
|---|---|---|---|---|---|---|
| **6I** | **Average velocity** | D2-Cre :: AAV-ChR2 (7/748) | OFF | 0.684±0.273 | Wilcoxon signed-rank test | ON-ChR2 vs. OFF-ChR2: *Z=47397*, *p*=$10^{-56}$ |
| | | | ON | -6.218±0.305 | | |
| | | D2-Cre :: AAV-eYFP (2/227) | OFF | 1.259±0.556 | Mann-Whitney U test | ON-ChR2 vs. ON-Control: *U=49134*, *p*=$10^{-22}$ |
| | | | ON | 0.189±0.520 | | |
| | | Vglut2-Cre :: AAV-ChR2 (7/778) | OFF | 0.142±0.245 | Wilcoxon signed-rank test | ON-ChR2 vs. OFF-ChR2: *Z=143970*, *p*=0.23 |
| | | | ON | 0.327±0.301 | | |
| | | Vglut2-Cre :: AAV-eYFP (2/158) | OFF | 0.850±0.530 | Mann-Whitney U test | ON-ChR2 vs. ON-Control: *U=58789*, *p*=0.19 |
| | | | ON | -0.064±0.526 | | |
| | | FoxP2-Cre :: AAV-ChR2 (6/711) | OFF | 0.566±0.179 | Wilcoxon signed-rank test | ON-ChR2 vs. OFF-ChR2: *Z=1395*, *p*=$10^{-114}$ |
| | | | ON | -15.535±0.139 | | |
| | | FoxP2-Cre :: AAV-eYFP (2/223) | OFF | 0.448±0.460 | Mann-Whitney U test | ON-ChR2 vs. ON-Control: *U=4559*, *p*=$10^{-100}$ |
| | | | ON | -0.163±0.444 | | |

*Table S7. Relates to Figure S1*

| Figure | Parameter | n (mice/neurons) | Data Type | Data Value (mean ± SEM) | Statistical test | significance level |
|---|---|---|---|---|---|---|
| **1E** | Proto Firing rate | (45/68) | labelled | 24,97 ± 1,38 | Mann-Whitney U test | U=3986, *p*=0,0065 |
| | | (46/152) | putative | 20,59 ± 0,81 | | |
| **1E** | Arky Firing rate | (22/23) | labelled | 3,532 ± 1,02 | Mann-Whitney U test | U=127,5, *p*=0,0066 |
| | | (19/21) | putative | 6,41 ± 0,83 | | |
| **1E** | Firing rate | (45/68) | Proto-GP labelled | 24,97 ± 1,38 | Mann-Whitney U test | U=42, *p*<0,0001 |
| | | (22/23) | Arky-GP labelled | 3,532 ± 1,02 | | |
| **1E** | Firing rate | (46/15) | Proto-GP putative | 20,59 ± 0,81 | Mann-Whitney U test | U=219, *p*<0,0001 |
| | | (19/21) | Arky-GP putative | 6,41 ± 0,83 | | |
| **1F** | Proto CV | (45/68) | labelled | 0,371 ± 0,02 | Mann-Whitney U test | U=5012, *p*=0,7217 |
| | | (46/152) | putative | 0,372 ± 0,01 | | |
| **1F** | Arky CV | (15/17) | labelled | 1,514 ± 0,21 | Mann-Whitney U test | U=126,5, *p*=0,1898 |
| | | (18/20) | putative | 1,16 ± 0,18 | | |
| **1F** | CV | (45/68) | Proto-GP labelled | 0,371 ± 0,02 | Mann-Whitney U test | U=56,50, *p*<0,0001 |
| | | (15/17) | Arky-GP labelled | 1,514 ± 0,21 | | |
| **1F** | CV | (46/152) | Proto-GP putative | 0,372 ± 0,01 | Mann-Whitney U test | U=191, *p*<0,0001 |
| | | (18/20) | Arky-GP putative | 1,16 ± 0,18 | | |
| **1I** | Proto Firing rate | (30/59) | labelled | 42.51 ± 3.48 | Mann-Whitney U test | U=2634, *p*=0.053 |
| | | (57/109) | putative | 48.57 ± 2.15 | | |
| | Arky Firing rate | (14/16) | labelled | 5.2 ± 1.27 | Mann-Whitney U test | U=493.5, *p*=0.221 |
| | | (46/75) | putative | 3.46 ± 0.54 | | |
| **1J** | Proto SDisi | (29/58) | labelled | 0.0109 ± 0.0034 | Mann-Whitney U test | U=3080, *p*=0.786 |
| | | (57/109) | putative | 0.0046 ± 0.0004 | | |
| | Arky SDisi | (9/9) | labelled | 0.0775 ± 0.0256 | Mann-Whitney U test | U=126, *p*=0.78 |
| | | (16/30) | putative | 0.395 ± 0.262 | | |

***Table S8. Relates to Figure S2***

| Figure | Parameter | n (mice/neurons) | Data Type | Data Value (mean ± SEM) | Statistical test | significance level |
|---|---|---|---|---|---|---|
| **2C** | D2-SPNs Firing rate | (4/8) | OFF | 0,018 ± 0,018 | Wilcoxon signed rank test | *p*=0,1250 |
| | | | ON | 5,62 ± 2,33 | | |
| **2C** | Proto Firing rate | (21/30) | OFF | 27,14 ± 2,15 | Wilcoxon signed rank test | *p*<0,0001 |
| | | | ON | 9,51 ± 1,79 | | |
| **2C** | Arky Firing rate | (13/16) | OFF | 2,54 ± 1,32 | Wilcoxon signed rank test | *p*=0,0002 |
| | | | ON | 10,17 ± 2,84 | | |
| **2C** | STN Firing rate | (5/10) | OFF | 4,84 ± 1,07 | Paired t test | t=3,444, *p*=0,0073 |
| | | | ON | 16,88 ± 4,03 | | |

***Table S9. Relates to Figure S3***

| Figure | Parameter | n (mice/neurons) | Data Type | Data Value (mean ± SEM) | Statistical test | significance level |
|---|---|---|---|---|---|---|
| **3A** | Put-D2-SPNs Firing rate | (7/40) | OFF | 0,061 ± 0,025 | Wilcoxon signed rank test | *p*<0,0001 |
| | | | ON | 25,92 ± 2,02 | | |
| **3A** | Put-Proto Firing rate | (22/91) | OFF | 19,39 ± 0,96 | Wilcoxon signed rank test | *p*<0,0001 |
| | | | ON | 0,908 ± 0,263 | | |
| **3A** | Put-Arky Firing rate | (9/10) | OFF | 5,74 ± 1,19 | Paired t test | t=6,681, *p*<0,0001 |
| | | | ON | 25,30 ± 3,24 | | |
| **3A** | Put-STN Firing rate | (7/53) | OFF | 4,07 ± 0,61 | Wilcoxon signed rank test | *p*<0,0001 |
| | | | ON | 15,56 ± 1,31 | | |
| **3B** | Put-D2-SPNs Firing rate | (7/25) | OFF | 0,066 ± 0,03 | Wilcoxon signed rank test | *p*<0,0001 |
| | | | ON | 9,34 ± 1,65 | | |
| **3B** | Put-Proto Firing rate | (41/180) | OFF | 26,73 ± 0,87 | Wilcoxon signed rank test | *p*<0,0001 |
| | | | ON | 12,57 ± 0,93 | | |
| **3B** | Put-Arky Firing rate | (20/22) | OFF | 2,96 ± 1,08 | Wilcoxon signed rank test | *p*=0,0002 |
| | | | ON | 9,76 ± 2,37 | | |
| **3B** | Put-STN Firing rate | (7/40) | OFF | 3,93 ± 0,78 | Wilcoxon signed rank test | *p*<0,0001 |
| | | | ON | 13,14 ± 2,04 | | |

***Table S10. Relates to Figure S4***

| Figure | Parameter | n (mice/neurons) | Data Type | Data Value (mean ± SEM) | Statistical test | significance level |
|---|---|---|---|---|---|---|
| **S4C** | Input recipient D2-MSN | (30/52) | % of Proto-putative AAV | 88.46 % | Fischer's exact test | *p*=0.4272 |
| | | (12/23) | % of Proto-putative Ai32 | 95.65 % | | |
| | Conductance | (25/46) | AAV | 19.86 ± 1.98 | Mann-Whitney U test | U=298, *p*=0.0058 |
| | | (10/22) | Ai32 | 41.87 ± 7.37 | | |
| **S4D** | Input recipient D2-MSN | (23/40) | % of Arky-putative AAV | 60% | Fischer's exact test | *p*=0.999 |
| | | (16/16) | % of Arky-putative Ai32 | 62.5% | | |
| | Conductance | (10/24) | AAV | 9.9 ± 2.73 | Mann-Whitney U test | U=84, *p*=0.182 |
| | | (7/10) | Ai32 | 20.09 ± 8.24 | | |
| **S4F** | Input recipient D2-MSN | (42/75) | % of Proto-putative | 90.66% | Fischer's exact test | *p*<0.0001 |
| | | (39/56) | % of Arky-putative | 60.71% | | |
| | Conductance | (35/68) | Proto-putative | 26.98 ± 2.98 | Mann-Whitney U test | U=565, *p*=0.0001 |
| | | (17/34) | Arky-putative | 12.9 ± 3.12 | | |

*Table S11. Relates to Figure S5*

| Figure | Parameter | n (mice/neurons) | Data Type | Data Value (mean ± SEM) | Statistical test | significance level |
|---|---|---|---|---|---|---|
| **5C (Top)** | STN Firing rate | (5/11) | OFF | 4,29 ± 1,27 | Paired t test | t=4,183, *p*=0,0019 |
| | | | ON | 22,36 ± 4,54 | | |
| **5C (Middle)** | Proto Firing rate | (29/39) | OFF | 22,39 ± 2,25 | Wilcoxon signed rank test | *p*<0,0001 |
| | | | ON | 75,63 ± 7,97 | | |
| **5C (Bottom)** | Arky Firing rate | (12/14) | OFF | 1,20 ± 0,66 | Wilcoxon signed rank test | *p*=0,0313 |
| | | | ON | 0,18 ± 0,099 | | |

***Table S12. Relates to Figure S6***

| Figure | Parameter | n (mice/neurons) | Data Type | Data Value (mean ± SEM) | Statistical test | significance level |
|---|---|---|---|---|---|---|
| **6A (Top)** | Put-STN Firing rate | (8/54) | OFF | 3,31 ± 0,53 | Wilcoxon signed rank test | *p*<0,0001 |
| | | | ON | 16,20 ± 1,52 | | |
| **6A (Middle)** | Put-Proto Firing rate | (21/61) | OFF | 22,05 ± 1,35 | Paired t test | t=14,23, *p*<0,0001 |
| | | | ON | 58,51 ± 3,43 | | |
| **6A (Bottom)** | Put-Arky Firing rate | (10/11) | OFF | 7,02 ± 1,19 | Paired t test | t=5.095, *p*=0,0005 |
| | | | ON | 1,36 ± 0,58 | | |
| **6B (Top)** | Put-STN Firing rate | (8/33) | OFF | 3,15 ± 0,53 | Wilcoxon signed rank test | *p*<0,0001 |
| | | | ON | 20,78 ± 2,36 | | |
| **6B (Middle)** | Put-Proto Firing rate | (28/122) | OFF | 27,91 ± 1,19 | Wilcoxon signed rank test | *p*<0,0001 |
| | | | ON | 82,88 ± 3,79 | | |
| **6B (Bottom)** | Put-Arky Firing rate | (14/15) | OFF | 5,71 ± 1,31 | Paired t test | t=3,048, *p*=0,0087 |
| | | | ON | 2,33 ± 0,64 | | |

***Table S13. Relates to Figure S7***

| Figure | Parameter | n (mice/neurons) | Data Type | Data Value (mean ± SEM) | Statistical test | significance level |
|---|---|---|---|---|---|---|
| **S7B** | Input recipient STN | (14/39) | % of Proto-putative | 94.87% | *Fischer exact test* | *p*=0.36 |
| | | (11/24) | % of Arky-putative | 87.5% | | |
| **S7C** | Conductance | (14/37) | Proto-putative | 3.74 ± 0.41 | Mann-Whitney U test | U=255, *p*=0.0306 |
| | | (10/21) | Arky-putative | 2.28 ± 0.34 | | |

*Table S14. Relates to Figure S8*

| Figure | Parameter | n (mice/neurons) | Data Type | Data Value (mean ± SEM) | Statistical test | significance level |
|---|---|---|---|---|---|---|
| **8F** | Arky Firing rate | (7/9) | OFF | 3,95 ± 1,39 | Friedman test with Dunn's multiple comparisons | Off *vs.* D2: Z=2.008, *p*=0.2677<br>Off *vs.* STN: Z=1.461, *p*=0.8648<br>Off *vs.* D2+STN: Z=2.008, *p*=0.2677<br>D2 *vs.* STN: Z=3.469, *p*=0.0031<br>D2 *vs.* D2+STN: Z=0.000, *p*>0.9999<br>STN *vs.* D2+STN: Z=3.469, *p*=0.0031 |
| | | | D2 | 23,56 ± 4,79 | | |
| | | | STN | 2,73 ± 2,73 | | |
| | | | D2 + STN | 37,94 ± 14,11 | | |
| **8G** | Arky Firing rate | (12/13) | OFF | 1,57 ± 0,94 | Friedman test with Dunn's multiple comparisons | Off *vs.* D2: Z=2.431, *p*=0.0904<br>Off *vs.* STN: Z=1.139, *p*>0.9999<br>Off *vs.* D2+STN: Z=1.747, *p*=0.4838<br>D2 *vs.* STN: Z=3.570, *p*=0.0021<br>D2 *vs.* D2+STN: Z=0.6836, *p*>0.9999<br>STN *vs.* D2+STN: Z=2.886, *p*=0.0234 |
| | | | D2 | 8,85 ± 2,38 | | |
| | | | STN | 2,23 ± 2,23 | | |
| | | | D2 + STN | 14,56 ± 2,23 | | |
